# Mapping the Peptide Interaction Fingerprint of the Behçet’s disease associated HLA-B*51

**DOI:** 10.1101/2025.11.26.690622

**Authors:** Sema Zeynep Yilmaz, Derman Basturk, Ahmet Gul, Burak Erman, Mert Gur

## Abstract

The strongest genetic risk factor for Behçet’s disease, a relapsing inflammatory disorder marked by recurrent mucocutaneous ulcers and uveitis, is an allele of the class I major histocompatibility complex (MHC-I) molecule, which presents intracellular peptides to CD8^+^ T cells. The molecular mechanisms linking the peptide preferences of this allele (HLA-B*51:01) to dysregulated immunity remain unclear, limiting efforts to design peptide-based modulators of antigen presentation. Here, we define HLA-B*51:01’s peptide selection rules by mapping the “interaction fingerprint” of 36 self-peptides using long-duration all-atom MD simulations. These uncovered a conserved hydrophobic–polar blueprint that is tuned by peptide length. High-speed atomic force microscopy rate *in silico* pulling experiments suggest a three-tier hierarchy of mechanical resilience: 9-mers resist the highest forces, 8-mers exhibit intermediate resistance, and 10/11-mers rupture most easily. Our comprehensive analysis provides an atomistic framework for understanding the molecular mechanisms underlying HLA-B*51:01 pathobiology and offers quantitative parameters to guide the design of therapeutic peptides or small molecules to modulate antigen presentation in Behçet’s disease.

**STATEMENT OF SIGNIFICANCE:** Behçet’s disease is strongly linked to HLA-B*51:01, a molecule that displays protein fragments to killer T cells, yet how this allele selects its peptides is poorly understood. Here, we combine all-atom equilibrium molecular dynamics simulations with steered MD pulling simulations that mimic high-speed atomic force microscopy experiments to map how 36 self-peptides of different lengths engage the HLA-B*51:01 groove. We uncover a conserved hydrophobic–polar interaction blueprint and show that 9-mers form the mechanically most resilient complexes, whereas shorter or longer peptides detach more easily under force. This length-tuned interaction “fingerprint” provides an atomistic framework for understanding HLA-B*51:01-driven immune dysregulation and guides the rational design of peptide-based or small-molecule modulators for Behçet’s disease.

## INTRODUCTION

Major histocompatibility complex (MHC) molecules play a critical role in the immune system by presenting antigenic peptides on the cell surface. These peptide–MHC complexes are recognized by T cells, which initiate an immune response if the peptide is derived from foreign proteins or abnormal self-proteins (1). In humans, MHC molecules are referred to as human leukocyte antigen (HLA). Based on their role in peptide presentation, HLA molecules are classified into class I and class II, with class I molecules further subclassified as HLA-A, HLA-B, and HLA-C, reflecting their genetic loci (2). HLA class I molecules present peptides derived from intracellular proteins, such as viral proteins, proteins from intracellular bacteria, self-peptides derived from self-proteins during their turnover or mutated self-proteins from cancer cells. By comparison, HLA class II molecules present peptides derived from extracellular proteins (e.g., proteins from extracellular bacteria or pathogens captured via phagocytosis) (3). Peptides of different lengths generated by proteasomal degradation of intracellular proteins are first translocated from the cytosol into the endoplasmic reticulum (ER) by the transporter associated with antigen processing (TAP), where they are further processed by aminopeptidases and loaded onto HLA class I molecules for presentation on the cell surface (4). The binding of a peptide stabilizes the HLA class I complex, allowing it to be transported via ER-derived vesicles through the Golgi apparatus to the cell surface (5,6). These peptide–HLA class I complexes are then displayed on the cell membrane for surveillance by CD8^+^ T cells (4). T cells scrutinize the entire HLA class I-peptide complex to discern any differences caused by altered self-proteins or foreign proteins. The T-cell receptor (TCR) binds primarily to the α1 and α2 helices of the HLA class I heavy chain, which together form the peptide-binding groove that houses the antigenic peptide (Figure 1) and interacts directly with both the peptide and these helices (7). Changes in the peptide sequence can alter molecular interactions at this interface, affecting how the TCR engages with the HLA class I-peptide complex, thereby modifying the TCR’s conformation and signaling responses, and enabling it to discern whether the peptide is self or foreign (8,9). When specific interactions occur between the TCR and peptide–HLA complexes displaying foreign or altered self-peptides, the TCR initiates the signaling cascade required for an effector immune response (10). Collectively, this process ensures the immune system recognizes and tolerates self-proteins while triggering an effector response upon detecting alterations in self-proteins or the presence of peptides from foreign organisms (11).

**Figure 1.**
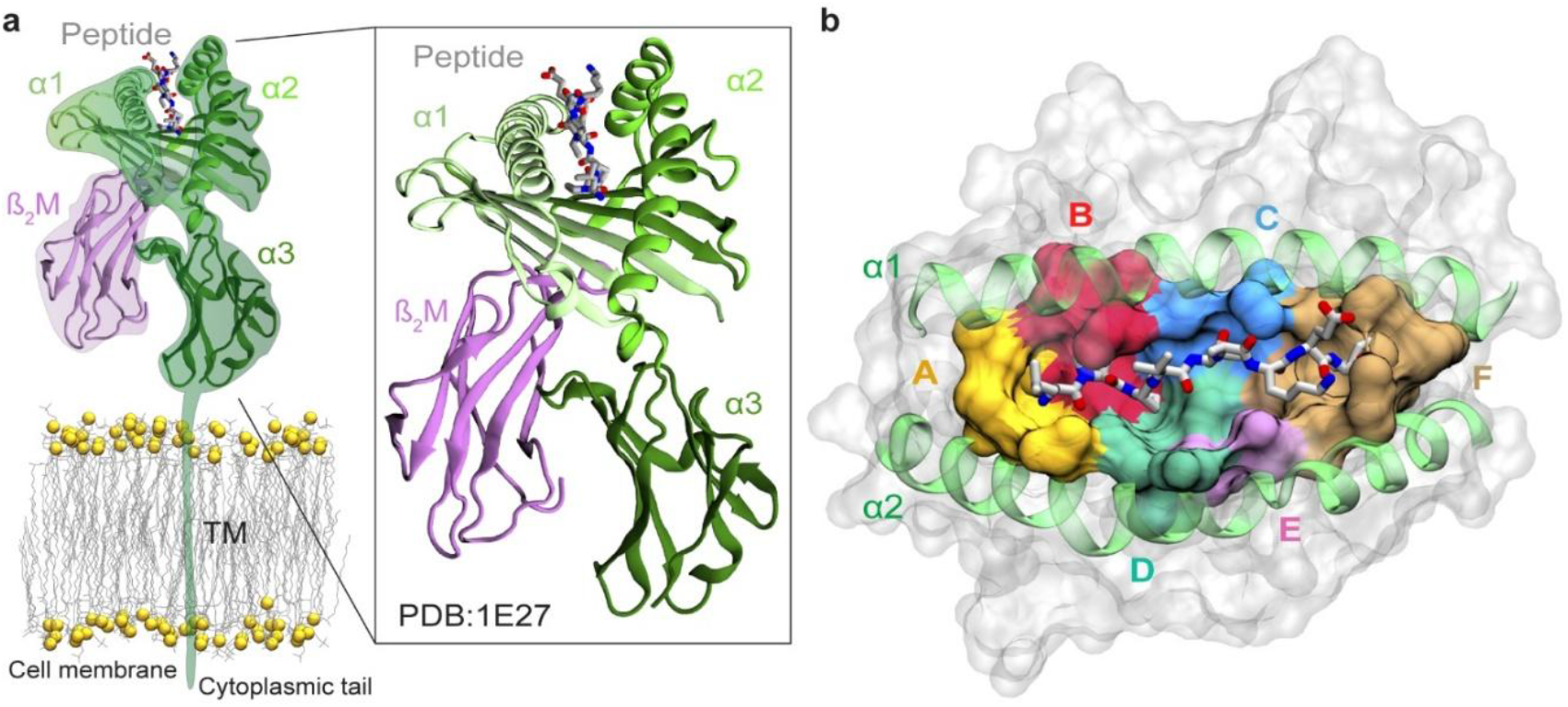
Structure of HLA-B*51:01 bound to an antigenic peptide. (a) Schematic representation of membrane-embedded HLA-B*51:01 presenting an antigenic peptide, LPPVVAKEI. The HLA-B*51:01 heavy chain (green) comprises five regions: the cytoplasmic tail, the transmembrane (TM) domain, and the α1, α2, and α3 domains. The light chain, β_2_M, is shown in purple. The peptide is depicted in stick representation, with carbon, nitrogen, and oxygen atoms colored gray, blue, and red, respectively. The heavy and light chains are shown in cartoon representation based on the experimentally determined structure (PDB: 1E27 (16)), whereas the transmembrane region, cytoplasmic tail, and lipid membrane are rendered schematically. (b) The A–F pockets within the peptide-binding groove of HLA-B*51:01 are displayed in surface representation. The flanking α-helices (α1 and α2 helices) are shown in green cartoon representation.

Due to their crucial role in the immune system, HLA molecules are strongly associated with various diseases. Across HLA class I molecules, disruptions in peptide selection or the stability of peptide– MHC complexes can misdirect CD8^+^ T-cell surveillance and precipitate inflammatory or autoimmune pathology. The HLA-B*51 allele, part of the B5 supertype of HLA-B, has been closely linked to Behçet’s disease, a multifactorial inflammatory condition of unknown etiology (12,13). Behçet’s disease is characterized by recurrent inflammatory episodes, leading to symptoms such as oral aphthous ulceration, genital ulceration, skin lesions, and uveitis. It has a strong genetic component, which is considered to be associated with immunological abnormalities (13). The prevalence of Behçet’s disease is especially high in populations along the historical Silk Road, which may correlate with the high frequency of the HLA-B*51 allele in healthy individuals from these regions (13,14). Furthermore, the HLA-B*51 allele is present in about 60% of patients with Behçet’s disease (15). This strong association with Behçet’s disease suggests that HLA-B*51 plays a significant role in the disease pathogenesis.

HLA-B*51:01 is a heterodimer composed of a heavy chain (α chain) non-covalently associated with a light chain (β2-microglobulin, β2M), a structure essential for antigen presentation (Figure 1a). The α chain includes a cytoplasmic tail, a transmembrane region, and extracellular domains (α1, α2, and α3), with α3 interacting directly with β_2_M (16). The peptide-binding groove of HLA class I, formed by the α1 and α2 domains, is flanked by α-helices and has a β-pleated sheet as its floor, forming six pockets (A–F) that collectively facilitate peptide binding and contribute to binding specificity (17,18) (Figure 1b). Residue assignments for pockets A–F and the roles of these pockets in peptide binding are described below based on general HLA class I definitions (17,18), with both the residue and pocket positions specifically mapped here onto the HLA-B*51:01 structure prior to detailed interaction analysis. The A pocket (residues M5, Y59, L163, W167, and H171) accommodates the peptide’s N-terminal residue. Adjacent to this, the B pocket (residues Y7, A24, V34, T45, N63, I66, and F67) stabilizes the peptide’s N-terminal region. Central pockets C (residues Y9, N70, T73, Y74) and D (residues Y99, Q155, Y159, and L160) anchor mid-peptide residues from opposite sides. The E pocket (residues T97, N114, E152, and L156) further supports peptide binding. The F pocket (residues N77, I80, A81, Y84, W95, Y116, Y123, T143, K146, and W147) anchors the peptide’s C-terminal residue. Together, these pockets facilitate precise peptide-HLA interactions, enabling effective antigen presentation and contributing to the peptide-binding specificity of HLA-B*51:01 (16,18).

HLA-B*51:01 typically binds peptides 8–11 amino acids in length, among which nonamers (9-mers) were the majority (61.7% of identified peptides), followed by octamers (8-mers) at 29.5%, with smaller proportions of decamers (10-mers) and undecamers (11-mers) at 4.8% and 3.0%, respectively (19,20). Studies of naturally processed peptides and synthetic peptides binding to HLA-B*51:01 have shown the presence of anchor residues at position 2 (P2) and the C-terminal position (PΩ) in peptides of 8–11 residues in length, as is common for HLA class I motifs (19-22). Residues at each position in the HLA-B*51:01 bound peptides were classified by their observed frequency of occurrence as *dominant* (>30%), *strong* (20–30%), or *preferred* (10–20%) (Table 1) (20). Asp and Leu frequently appear at P1 (strong in 8-mers to 10-mers, preferred in 11-mers), while Pro (dominant) and Ala (dominant in 9-mers, strong in 8-mers and 10-mers, and preferred in 11-mers) are the most common at P2. P3 often hosts aliphatic or aromatic residues (preferred to dominant), and Val or Ile (mostly dominant) occupy PΩ (20). HLA-B*51:01 bound peptides show a potential correlation between the first two positions P1 and P2: when Pro occupies P2, about 80% of peptides carry an aliphatic or aromatic residue at P1, whereas when Ala is present at P2, Asp is found at P1 in roughly 55% of cases (20).

**Table 1.** Position-specific frequencies of the most common residue types in HLA-B*51:01 binding peptides. The table was adapted based on Figure 2 in Guasp et al. (20), which was obtained by investigating 1,620 peptide sequences, identified by mass spectrometry from HLA-B*51:01 molecules isolated from 721,221 transfectant cells, with subsequent analysis of residue frequencies at each peptide position to define the binding motif.

|  | P1 | P2 | P3 | P4 | P5 | P6 | P7 | P8 |
| --- | --- | --- | --- | --- | --- | --- | --- | --- |
| Dominant |  | P |  |  |  |  |  | I<br>V |
| Strong | D<br>L | A | Y |  |  | P |  | L |
| Preferred |  |  | P<br>F<br>L | L<br>V | V<br>L<br>A | L<br>S | T<br>L<br>K |  |

|  | P1 | P2 | P3 | P4 | P5 | P6 | P7 | P8 | P9 | P10 |
| --- | --- | --- | --- | --- | --- | --- | --- | --- | --- | --- |
| Dominant |  | P | Y |  |  |  |  |  |  | I<br>V |
| Strong | L<br>D | A |  |  |  |  |  |  | T | L |
| Preferred | Y |  | L<br>F | P<br>S<br>E | G<br>S | G<br>P<br>K | P | L<br>T<br>S<br>V<br>I | V |  |

|  | P1 | P2 | P3 | P4 | P5 | P6 | P7 | P8 | P9 |
| --- | --- | --- | --- | --- | --- | --- | --- | --- | --- |
| Dominant |  | P<br>A |  |  |  |  |  |  | V<br>I |
| Strong | D<br>L |  | Y |  |  |  |  | T |  |
| Preferred |  |  | L<br>F | P<br>E | P | V | P<br>L<br>S | S<br>L |  |

|  | P1 | P2 | P3 | P4 | P5 | P6 | P7 | P8 | P9 | P10 | P11 |
| --- | --- | --- | --- | --- | --- | --- | --- | --- | --- | --- | --- |
| Dominant |  | P | Y |  |  |  |  |  |  | T | I<br>V |
| Strong |  |  |  | P |  | S<br>P |  |  | L |  |  |
| Preferred | L<br>Y<br>I<br>D | A | F | E<br>D<br>G<br>A | A<br>S<br>L | G<br>D | A<br>K<br>G | T | R | V |  |

Two primary mechanisms have been suggested for the pathogenic role of HLA-B*51 in Behçet’s disease: (i) altered peptide presentation, which may lead to abnormal activation of CD8^+^ cytotoxic T cells and natural killer (NK) cells (23-25) and (ii) low-affinity peptide binding by HLA-B*51 may cause slow assembly and misfolding of HLA-B*51, triggering an endoplasmic reticulum stress response and an unfolded protein response that leads to inflammation (19,23). For both potential mechanisms, it is imperative to gain a molecular understanding of how HLA-B*51 interacts with its self-peptides and how these interactions affect its structure and dynamics. Such insights will not only elucidate the structural dynamics of HLA-B*51 under varying binding conditions but also potentially facilitate the development of novel therapeutics targeting HLA-B*51 by identifying and characterizing specific peptide-binding motifs, offering new strategies for treating Behçet’s disease.

However, although thousands of peptides can bind to the antigen-binding groove of HLA-B*51:01, there is limited information regarding these peptides, including their binding poses, affinities, and effects on the structural dynamics of HLA-B*51:01. Experimentally obtained structures for peptides in complex with HLA-B*51:01 exist only for viral peptides TAFTIPSI (16,26) and LPPVVAKEI (16). Furthermore, bound conformations of self-peptides in complex with HLA-B*51:01, obtained through all-atom molecular dynamics (MD) simulations, are available only for four peptides: YAYDGKDYI, LPRSTVINI, IPYQDLPHL, which were reported in our previous study (27), and AAAAAIFVI (28). Consequently, atomic-level structural insights into peptide–HLA-B*51:01 interactions are currently restricted to just these six peptides.

To address the current lack of structural and dynamic understanding of peptide–HLA-B*51:01 interactions, we conducted a comprehensive systematic analysis on a wide range of peptides reported to bind to HLA-B*51:01, selected from the Immune Epitope Database (29). Our selection criteria focused on peptides ranging from 8 to 11 amino acids, particularly those containing strong and dominant residues for HLA-B*51:01 binding (Table 1), yielding a total of 43 peptides. We utilized the binding affinities from NetMHCpan 4.1 (30) to narrow the selection of 43 peptides to 33, focusing on those with the highest predicted affinity for HLA-B*51:01 (Table S1). These peptides were subjected to 4.7 μs of conventional MD (cMD) simulations in the presence of HLA-B*51:01 to examine their binding poses and interactions. Combining simulations of 33 newly selected self-peptides with our three previously (27) simulated self-peptides (36 in total), we generated an interaction map highlighting key motifs involved in HLA-B*51:01 self-peptide binding. Furthermore, we conducted a total of 2.52 μs of *in silico* pulling experiments using steered MD (SMD) at loading rates comparable to those employed in high-speed atomic force microscopy (HS-AFM) experiments. These rates are one to two orders of magnitude lower than standard SMD loading rates, significantly minimizing entropy generation and thus providing more accurate estimates. Our SMD simulations focused on eight peptides, comprising the two highest-scoring candidates from each peptide length (8-mer to 11-mer) identified by NetMHCpan 4.1 (30), and provided rupture forces as well as unique atomic-level insights into their binding/unbinding processes with HLA-B*51:01.

## METHODS

### MD simulations

The X-ray structure of HLA-B*51:01 bound to its HIV immunodominant epitope (2.20 Å resolution; PDB ID: 1E27 (16)) served as the starting model. Each complex was embedded in a rectangular TIP3P water box that extended 25 Å beyond the solute in ±x and ±y (total 50 Å) and 15 Å in ±z, then charge-neutralized and brought to 150 mM NaCl. The fully solvated systems comprised ∼100,000 atoms. All simulations were carried out in NAMD (31) with the CHARMM36 force field (32) using a 2 fs integration step. The temperature was kept at 310 K with Langevin dynamics (damping coefficient 1 ps−^1^), and the pressure at 1 atm with a Langevin Nosé–Hoover barostat (oscillation period 100 fs, damping 50 fs). Van der Waals interactions were truncated at 12 Å, and long-range electrostatics were treated with particle-mesh Ewald (PME). After 10,000 steps of energy minimization, each system underwent a 1 ns equilibration in which all protein atoms were harmonically restrained. A second 10,000 step minimization was followed by 2 ns of equilibration with a 2 kcal mol/Å^2^ restraint on backbone atoms only. Unrestrained production trajectories of 100 ns were then recorded for every HLA-B*51:01–peptide complex (Table S2), saving frames every 50 ps for analysis.

### SMD simulations

For SMD simulations, the starting conformations were taken from the ends of cMD trajectories. Systems were solvated having a 40 Å cushion along the pulling direction to create enough space for unbinding simulations and a 15 Å cushion in all other directions. Ions were added to neutralize the system, and NaCl concentration was set to 150 mM. To allow the systems to reach equilibrium, we conducted 10,000 minimization steps and subsequently performed 1 ns equilibration simulations with the protein restrained. Then, we performed an additional 1 ns of equilibration for each HLA-B*51:01-peptide complex without any restraint. The final equilibrated systems were used as starting systems for SMD simulations. Each SMD simulation was performed until the rupture event was observed for the peptides (Table S3). In the SMD simulations, HLA-B*51:01 was fixed at its 97^th^ residue backbone atoms, and a dummy atom was attached to the fifth residue of the peptide via a virtual spring (k = 0.1439 kcal/mol/Å^2^), a value previously used to model HS-AFM springs *in silico* (33). This dummy atom was then pulled with a constant velocity of 0.1 Å/ns along a vector pointing from the center of mass of fixed atoms to the center of mass of steered atoms.

SMD calculations started from the final snapshots of the corresponding cMD runs. The complexes were resolvated with a 40 Å water cushion along the intended pulling axis and 15 Å in the remaining directions. Systems were neutralized and adjusted to 150 mM NaCl. Following 10,000 steps of minimization, the systems were equilibrated for 1 ns with protein backbone atoms restrained, and for an additional 1 ns without restraints. The pulling vector was defined from the center of mass of the fixed group (backboned atoms of residue T97) to that of the steered group (backbone atoms of P5), and the dummy atom was translated along this vector at a constant velocity until peptide dissociation (rupture) was observed.

### PCA

Trajectory snapshots were subjected to PCA as described previously (34-36). The Cartesian coordinates of all peptide C^α^ atoms were assembled into a configuration matrix, mean-centered, and used to build the covariance matrix. Diagonalization yields 3N-6 non-trivial eigenvectors (principal components, PCs), where N is the number of atoms included in the analysis. PC1 captures the largest variance (the dominant collective motion), PC2 the next largest, and so forth. All PCA calculations were performed with custom scripts written in VMD (Tcl) and MATLAB. The resulting trajectories were projected onto PC1 and PC2 to map the conformational landscape of peptide binding (Figures S1-S3).

## RESULTS AND DISCUSSION

### Generation and filtering of a peptide library for HLA-B*51:01 binding prediction

As a first step, the Immune Epitope Database (29) was used to generate a library of peptides that interact with HLA-B*51:01. This library comprised approximately 15.9k members in 2019 when we initiated this project, with peptide lengths ranging from 7 to 25 amino acids. Particularly, 94.4% of the library consisted of peptides 8-11 amino acids in length. As of 2025, the data has not changed considerably, having ∼16.5k members. As a next step, these ∼15.9 k peptides were filtered such that, for each peptide, all positions annotated as “Dominant” in Table 1 (20) were occupied by one of the corresponding dominant residues for the given peptide length (for an 8-mer, for example, P2 had to be P and P8 either I or V), resulting in a smaller subset of 6476 peptides; specifically, 1422 8-mers, 4972 9-mers, 75 10-mers, and seven 11-mers. After the second round of filtering, in which we further restricted this subset by additionally requiring that all positions annotated as “Strong” in Table 1 (20) were occupied by one of the corresponding strong residues (unless they are already occupied by a “Dominant” residue), while still fulfilling all dominant-position requirements of the first filtering step, a total of 36 peptides were obtained: 17 8-mers, 12 9-mers, and seven 10-mers. Because the second round of filtering didn’t yield any 11-mers, we kept the seven 11-mers resulting from the first filtering step. Thus, a library comprising 43 peptides was constructed (Table S1). We then utilized NetMHCpan 4.1 (30), a web tool that predicts peptide-MHC binding affinities using artificial neural networks trained on experimental data, to estimate the binding affinity of these 43 peptides to HLA-B*51:01 (Table S1). Among the 43 peptides, we selected all 9-mers, 10-mers, and 11-mers, along with the seven 8-mers that NetMHCpan 4.1 (30) predicted to have the highest binding affinity, thereby narrowing our focus to the top 33 peptides.

Structural models were constructed for each of the 33 peptides in complex with HLA-B*51:01, using the crystal structure of LPPVVAKEI bound to HLA-B*51:01 (PDB: 1E27 (16)) as a template. In the HLA-B*51:01 bound 9-mers models, the backbone coordinates and side chain rotations observed in LPPVVAKEI (set I) were maintained. For each of the 8-mers, on the other hand, two models were built using either P2-P9 or P1-P8 of LPPVVAKEI as templates (set II-a and set II-b, respectively). For each of the 10-mers, two structural models were constructed by adding a single residue either at the N- or C-terminal of the template peptide (set III-a and set III-b, respectively). For the 11-mers, the reference peptide was extended at both the N- and C-terminal ends by a single residue to build our models (set IV). 100 ns of cMD simulations were performed for each of these 47 structural models (totaling 4.7 μs) after performing two cycles of minimization and equilibration (Table S2). Along with the 47 newly generated cMD trajectories for 33 peptides, six cMD simulations (totaling 2.4 μs ) from our previously published study (27) on three self-peptide HLA-B*51:01 complexes were incorporated into the analysis, thereby increasing the total number of investigated self-peptides to 36. For these three previously studied self-peptides, the prior simulations included two 150 ns cMD runs for LPRSTVINI and IPYQDLPHL, and two 900 ns runs for YAYDGKDYI (totaling 2.4 μs).

All cMD trajectories were aligned to the HLA-B*51:01 crystal structure (PDB: 1E27(16)) by superposition of the C^α^ atoms from the HLA-B*51:01’s α1 and α2 helices. This procedure removes all rigid-body translations and rotations of the heavy chain, thereby isolating the peptide’s motion with respect to the binding groove. To quantify that motion, root-mean-square deviation (RMSD) values were computed for the peptide C^α^ atoms, capturing both global displacement and any internal conformational rearrangements. In nearly every simulation, the RMSD traces reached a clear plateau (Figures S4-S7), indicating that most peptides optimized into stable binding poses in their MD simulation runs. For each peptide, the binding pose in the MD run with the lower plateau RMSD was selected as the representative binding mode for downstream analyses, as it exhibited the smallest deviation from its starting conformation. To characterize dominant peptide conformations, we performed principal component analysis (PCA) (34-36) on the peptide C^α^ coordinates. Snapshots from each cMD trajectory were then projected onto the first two PCs, PC1 and PC2 (Figures 2 and S1-S3). The highest-density region in the resulting PC1–PC2 landscape, corresponding to the most frequently sampled conformational basin, was selected as the representative bound pose for that trajectory (Figure 2 and S8-S9).

**Figure 2.**
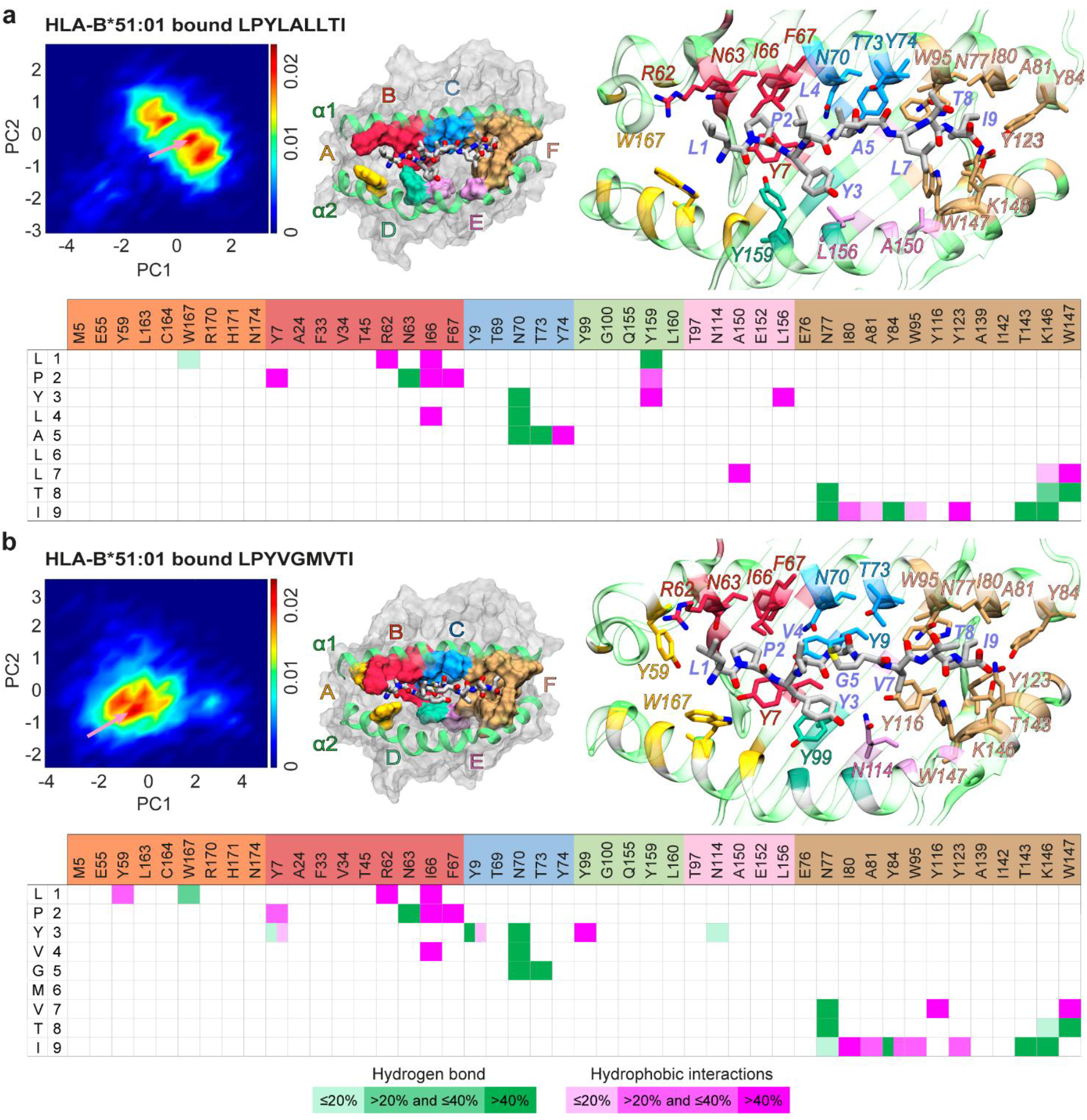
Conformational landscapes and interaction fingerprints of LPYLALLTI and LPYVGMVTI bound to HLA-B*51:01. (a) shows the LPYLALLTI–HLA-B*51:01 complex, and (b) depicts the LPYVGMVTI–HLA-B*51:01 complex. In each case, the top-left subpanel presents a two-dimensional conformational density plot obtained by projecting the ensemble of peptide conformations onto PC1 and PC2, upon structural alignment of the MD conformations via the HLA-B*51:01 binding groove. Next to it, the central subpanel illustrates a representative bound peptide conformation from the most densely populated (pink arrow, top-left panel) binding mode, shown in licorice representation, together with the binding pockets (A–F) of HLA-B*51:01 highlighted on the molecular surface. On the top-right, residue-level interaction networks for this representative pose are shown, with peptide carbon atoms colored gray and HLA carbon atoms colored according to their associated binding pocket. The lower subpanel provides an interaction matrix summarizing the peptide–HLA contacts across five selected peptide conformations within the most populated conformational basin. Hydrophobic interactions are indicated in magenta and hydrogen bonds in green, with shading intensity corresponding to interaction frequency across the selected five peptide conformations.

### Detailed interaction analysis reveals stable hydrophobic clusters and complementary polar interactions in peptide–HLA-B*51:01 complexes

For each binding mode, five representative poses were selected for detailed analysis in the Discovery Studio Visualizer (37). Across these poses, the peptides were engaged in hydrogen bonds, salt bridges, electrostatic interactions (π-cation/π-anion), π-sulfur interactions, and hydrophobic interactions (alkyl, π-alkyl, amide-π stacked/π-π stacked and π-σ) (Figure 2 and Tables S4–S8). For each peptide, interaction frequencies were classified as low (20%; observed in one conformation of the selected five representative conformations), medium (40%; observed in two conformations), or high (> 40%; observed in at least three conformations). LPYLALLTI and LPYVGMVTI, which showed the highest two binding affinity predictions by NetMHCpan 4.1 (30), each formed two well-defined hydrophobic clusters at the peptide–HLA-B*51:01 interface. In LPYLALLTI, hydrophobic-cluster 1 involves peptide residues L1, P2, Y3 and L4 together with HLA-B*51:01 residues Y7, R62*, I66, F67, L156, and Y159, whereas hydrophobic-cluster 2 comprises peptide residues L7 and I9 with HLA-B*51:01 residues I80, A81, W95, Y123, K146, W147, and A150* (Figures 3). In LPYVGMVTI, hydrophobic-cluster 1 involves peptide residues L1, P2, Y3, and V4 together with HLA-B*51:01 residues Y7, Y9, Y59, R62*, I66, F67, and Y99 while hydrophobic-cluster 2 comprises peptide residues V7 and I9 with HLA-B*51:01 residues I80, A81, Y84, W95, Y116, Y123, W147. These persistent hydrophobic interaction clusters are complemented by an extensive hydrogen bonding network. A comparable two-cluster hydrophobic arrangement, or a merged single large cluster, often reinforced by hydrogen bonds and, in 67% of cases additionally by one to three salt bridges, was observed for all peptides, although the composition and persistence of the clusters varied. (Figures 3 and S10-S11 and Tables S4–S8).

**Figure 3.**
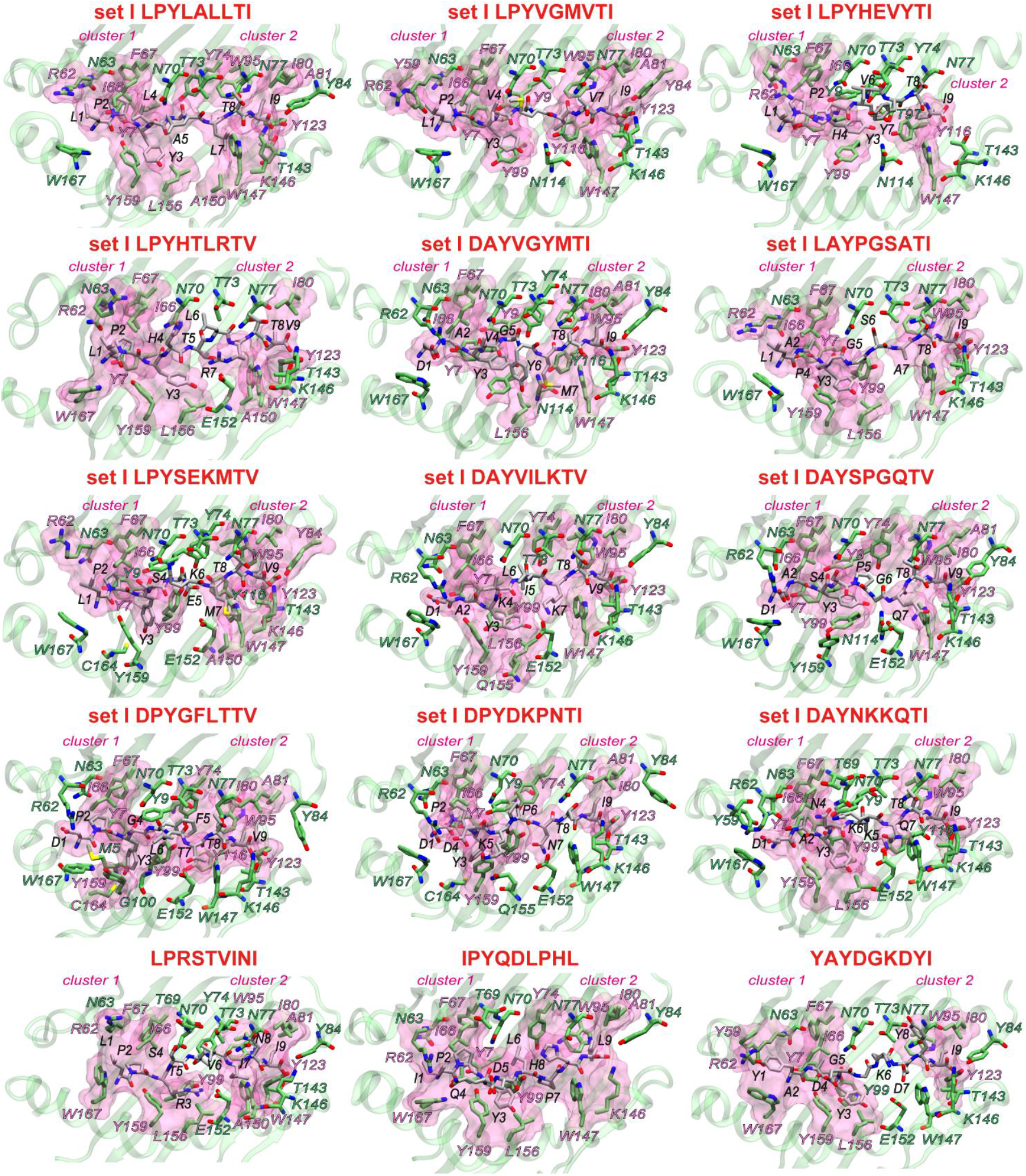
Two hydrophobic clusters consistently emerge at the interface of HLA-B*51:01 and 9-mers, stabilized by a dense hydrogen bonding network. Hydrophobic clusters are depicted as transparent magenta surface balloons. Peptide and interacting HLA-B*51:01 residues are shown in licorice representation, with the overall HLA-B*51:01 binding groove rendered in green cartoon representation. Peptide residues are labeled in black. HLA-B*51:01 residues forming hydrophobic contacts with the peptide are labeled in magenta, whereas residues involved in other non-covalent interactions (hydrogen bonds, salt bridges, electrostatic, or π-sulfur interactions) are labeled in green. The top four rows show peptides from the filtered list (set I) generated in this study, while the bottom row shows peptides studied in our earlier work (27).

### Interaction fingerprints reveal length-dependent engagement patterns and structural adaptability in peptide–HLA-B*51:01 complexes

Collective mapping of peptide–HLA-B*51:01 contacts reveals a coherent yet length-tunable interaction fingerprint across pockets A–F of the binding groove (Figure 4). Independent of peptide length, the most frequently observed interactions occur consistently with residues W167 in pocket A; Y7, R62, N63, I66, and F67 in pocket B; Y9, N70, T73, and Y74 in pocket C; Y99 and Y159 in pocket D; E152 in pocket E; and N77, I80, A81, Y84, W95, Y116, Y123, T143, K146, and W147 in pocket F.Notably, pocket F consistently serves as the dominant anchor for the C-terminal residue (PΩ) across all peptide lengths.

**Figure 4.**
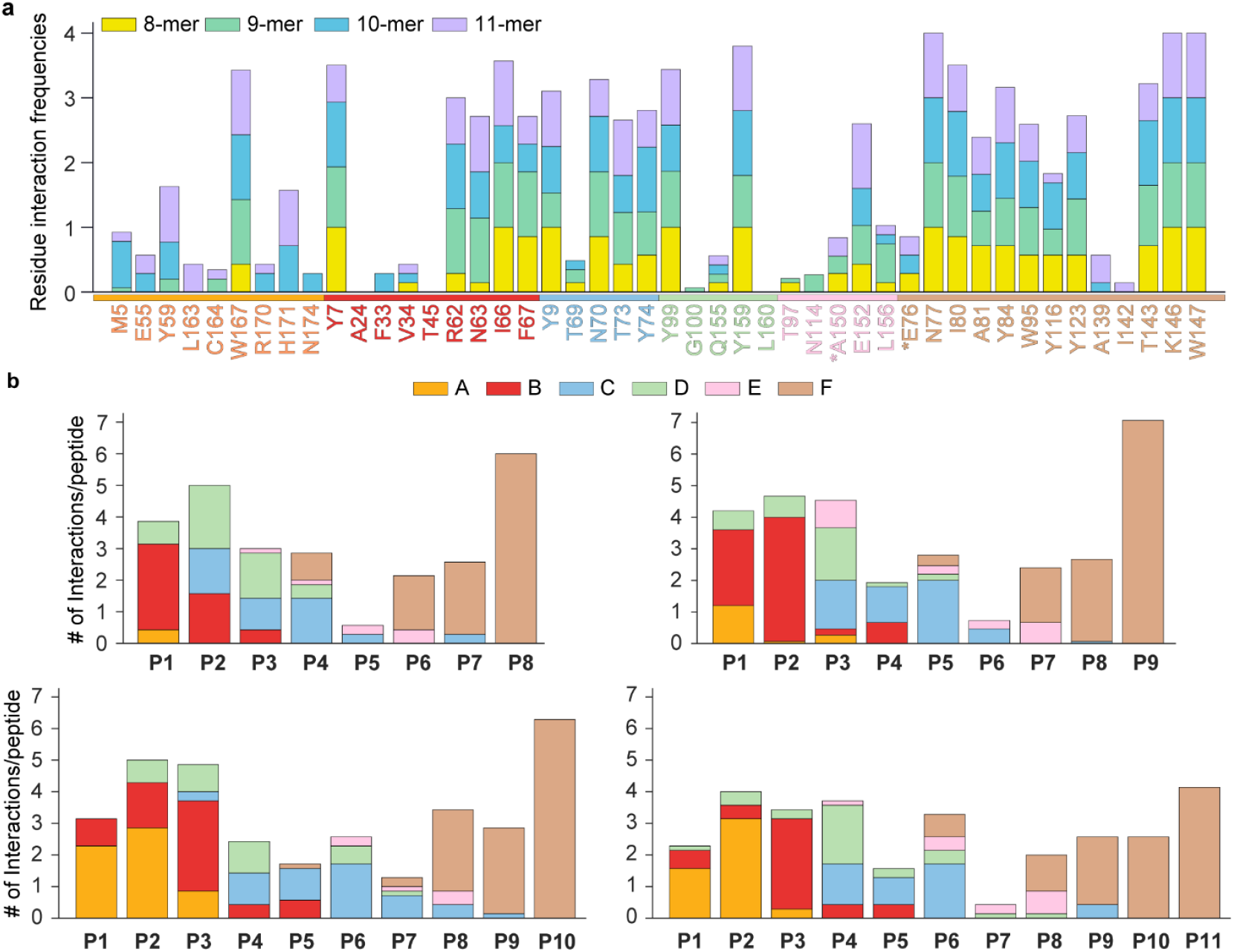
Length-dependent interaction patterns of peptides bound to HLA-B*51:01. (a) quantifies, for every HLA-B*51:01 residue that lines the groove, the mean number of contacts established by peptides of four different lengths (8-, 9-, 10-, and 11-mers). For a given length, contact counts from all peptides were pooled and then divided by the number of peptides in that set, so the resulting stacked bars report the average interactions contributed by each residue and directly expose length-specific preferences. Bar charts are separately provided for each peptide length in Figure S12. (b) turns the viewpoint from the HLA molecule to the peptide. Each of the four subplots corresponds to one peptide length and displays how many contacts, again normalized per peptide, originate from pockets A through F (color-coded) at each peptide position (P1–P11). Together, the two panels reveal how both residue usage within HLA-B*51:01 and positional engagement along the peptide vary as a function of peptide length.

Superimposed on these conserved interactions are length-dependent differences that alter anchoring patterns, particularly at N-terminal and central pockets. For example, 8-mer peptides exhibit additional interactions with residues V34 (pocket B), T69 (C), Q155 (D), T97, A150*, and L156 (E), and E76* (F), reflecting their preference for mid-groove stabilization. At position P1, 8-mers primarily interact with pocket B, while at position P2, they interact more broadly with pockets A, B, and D.

In contrast, 9-mers engage additional residues such as M5, Y59, and C164* (pocket A), T69 (C), G100* and Q155 (D), T97, N114, A150*, and L156 (E). These peptides exhibit the canonical class I anchoring motif (P2 and PΩ anchoring)(19-22), notably characterized by strong anchoring at P2 in pocket B and dual-pocket interactions (A and B) at P1.

Longer peptides, such as 10-mers and 11-mers, further modulate interaction profiles. 10-mers interact additionally with residues M5, E55*, Y59, R170*, H171, and N174* (pocket A); F33* and V34 (B); T69 (C); Q155 (D); L156 (E); and E76* and A139* (F). These peptides demonstrate enhanced interactions at pocket A for positions P1 and P2, signifying a shift in the N-terminal anchoring region with increasing peptide length. Similarly, 11-mers expand their interactions further with residues M5, E55*, Y59, L163, C164*, R170*, and H171 (A); V34* (B); Q155 (D); A150* and L156 (E); and E76*, A139*,and I142* (F), predominantly anchoring at pocket A for P1 and reengaging pocket B at P2.

Across all peptide lengths, residues at positions P4–P6 consistently contribute to binding stability through interaction with pocket C. Collectively, these observations illustrate the structural adaptability of HLA-B*51:01, highlighting its capability to accommodate peptides of varying lengths by dynamically adjusting its interaction patterns, especially at the N-terminal and central positions. These molecular insights provide a deeper understanding of peptide selection mechanisms pertinent to antigen presentation pathways and Behçet’s disease pathogenesis.

### Exploring the binding strength of peptides known to bind to HLA-B*51:01

We conducted *in-silico* pulling experiments using all-atom SMD simulations to quantify the binding strength of peptides to HLA-B*51:01 under externally applied forces. In an SMD simulation, a dummy atom is connected by a harmonic spring to a user-defined set of steered atoms; the dummy atom is then translated at constant velocity, thereby loading the system through the spring. To mimic the loading regime of HS-AFM measurements (33), we used a spring constant of 0.1438 kcal/mol/Å^2^ (equivalent to 10 pN/Å) and a pulling speed of 0.1 Å/ns. These parameters produce a soft-spring protocol, in contrast to the stiff-spring approximation commonly employed in SMD, where a much larger force constant constrains the steered atoms to track the dummy atom almost exactly. A soft spring allows the force to build up gradually, closely reproducing the HS-AFM experimental condition in which the cantilever bends slowly until the complex ruptures; hence, the data generated here are directly comparable to the experiment.

In the SMD simulations, the backbone atoms of the fifth residue of each peptide were selected as the steered atoms, while the backbone atoms of T97 of HLA-B*51:01 were restrained to represent immobilization of the HLA (Figure 5a). SMD simulations were performed for the two peptides with the highest NetMHCpan 4.1 (30) scores at each peptide length. The simulations showed that peptide length exerts a dominant influence on mechanical stability: The 9-mer peptides LPYLALLTI and LPYVGMVTI resisted the highest forces (≈ 350–380 pN) and required the greatest work (≈ 100– 115 kcal/mol) for extraction. The 8-mer peptides LPYIFPNI and LPYPDPAI display intermediate rupture forces (≈ 250–260 pN) and work values (≈ 50 kcal/mol) (Figure 5c and S13). The 10-mer peptides DPYSFGRTTI and DAYNSLSSTV and 11-mer peptides IPYDAKTIQTI and YPYNQPKRNTI detach at markedly lower forces (≈ 100–200 pN) and with less work (< 50 kcal/mol). These differences are visualized in Figure 5b, which plots rupture force versus work, and in the individual force– extension traces of Figure 5c, where the peak force for each peptide is indicated numerically (in pN). The steep extended plateaus for the 9-mers indicate that these peptides maintain simultaneous anchoring in multiple HLA pockets throughout most of the pulling coordinate, whereas the shorter (8-mer) and longer (10/11-mer) peptides lose critical contacts earlier, leading to lower force maxima and less work accumulation.

**Figure 5.**
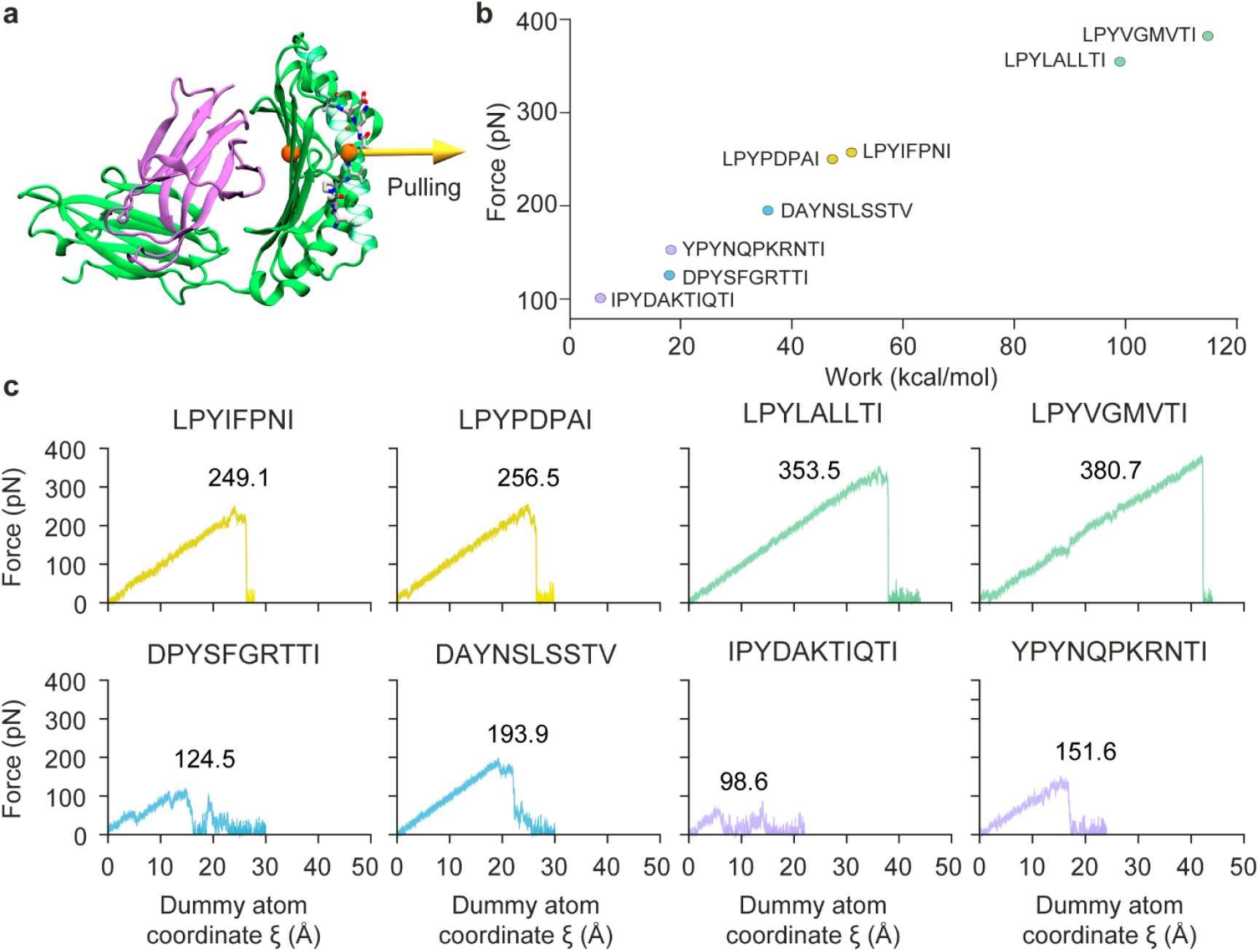
9-mers exhibit the highest binding strengths to HLA-B*51:01. (a) In the *in-silico* pulling set-up, the backbone atoms of residue 97 of HLA-B*51:01 were fixed, while the backbone atoms of the fifth of each peptide were steered. The yellow arrow denotes the SMD pulling vector, which served as the reaction coordinate. (b) Scatter plot comparing the rupture force (ordinate) and the mechanical work (abscissa) recorded in SMD simulations for each peptide. (c) Force–extension profiles for all peptides; the peak rupture force (pN) is printed within each panel.

## CONCLUSION

This work set out to provide an atomic-level description of how HLA-B*51:01 selects, stabilizes, and ultimately presents self-peptides of different lengths, a question that is central both to basic antigen-presentation biology and to its potential role in the pathogenesis of Behçet’s disease. By combining 7.1 µs of cMD simulations on 36 peptide–HLA complexes with 2.5 µs of HS-AFM rate constant-velocity SMD simulations, we uncovered a coherent yet length-tunable interaction blueprint for pockets A–F and linked that blueprint to the mechanical resilience of the complex under load.

Our equilibrium simulations confirm the canonical P2/PΩ anchoring motif for 9-mers, but also reveal systematic shifts when the peptide is one residue shorter or longer. 8-mer draws additional stabilization from the mid-groove (B–D pockets), whereas 10-mers and 11-mers advance their N-terminal anchor toward pocket A to compensate for steric strain introduced potentially by backbone bulging. These length-dependent adaptations rationalize why 9-mers dominate the naturally processed immunopeptidome of HLA-B*51:01.

Sequence-to-structure mapping highlights two persistent hydrophobic clusters that knit together the peptide and HLA surfaces and are complemented by a network of hydrogen bonds and salt bridges. Critically, residues N63 and F67, positions that differ between the disease-associated HLA-B*51 and the non-associated HLA-B*52, emerge as key contributors to the first hydrophobic cluster for 9-mers but are bypassed or only transiently engaged by 10-/11-mers. This observation provides a structural rationale for the long-standing hypothesis that subtle alterations in pocket B help dictate disease risk.

The scatter plot in Figure 5b groups the peptides into three clearly separated regimes of mechanical stability: high-strength 9-mers, mid-strength 8-mers, and low-strength 10-/11-mers. This trend underscores the structural match between the canonical 9-mer length and the dimensions of the HLA-B*51:01 groove, which enables optimal engagement of both primary (A/F) and secondary (B–E) anchor pockets. Shorter 8-mers cannot fully occupy the F-pocket, reducing their resistance to force, while longer peptides bulge out of the groove, likely incurring steric strain that weakens pocket contacts and leads to premature rupture. The force traces in Figure 5c corroborate this interpretation: 9-mers exhibit long, linear force ramps ending in abrupt drops characteristic of cooperative unbinding, whereas 10/11-mers display noisy, multi-step profiles indicative of sequential contact loss. Collectively, the SMD results provide a mechanistic rationale for the experimental observation (20) that 9-mer dominate the HLA-B*51:01 immunopeptidome and suggest that mechanical robustness contributes to efficient antigen presentation.

In sum, our integrated computational approach delineates how peptide length, anchor-pocket complementarity, and cooperative hydrophobic clustering synergize to govern both thermodynamic affinity and mechanical durability in HLA-B*51:01. These insights deliver a quantitative framework for interpreting allele-specific disease associations and will aid the rational design of peptide mimetics or small molecules aimed at modulating HLA-B*51:01 presentation in Behçet’s disease.

## AUTHOR CONTRIBUTIONS

Conceptualization: M.G., A.G., and B.E.; Methodology: S.Z.Y. and M.G.; Investigation: S.Z.Y. and M.G.; Analysis and Visualization: S.Z.Y., D.B., and M.G. Writing−original draft: S.Z.Y., D.B., and M.G. Writing−review and editing: S.Z.Y., A.G., B.E., and M.G.

## ACKNOWLEDGEMENTS

M.G. acknowledges support from the Scientific and Technological Research Council of Türkiye (TÜBİTAK) under grant no. 119Z553, and from his start-up funds provided by the University of Pittsburgh School of Medicine.

