## Supplementary material for "Mapping the Peptide Interaction Fingerprint of the Behçet’s disease associated HLA-B*51": Suplementary Information

### Supplemental Information

#### Supplementary Figures

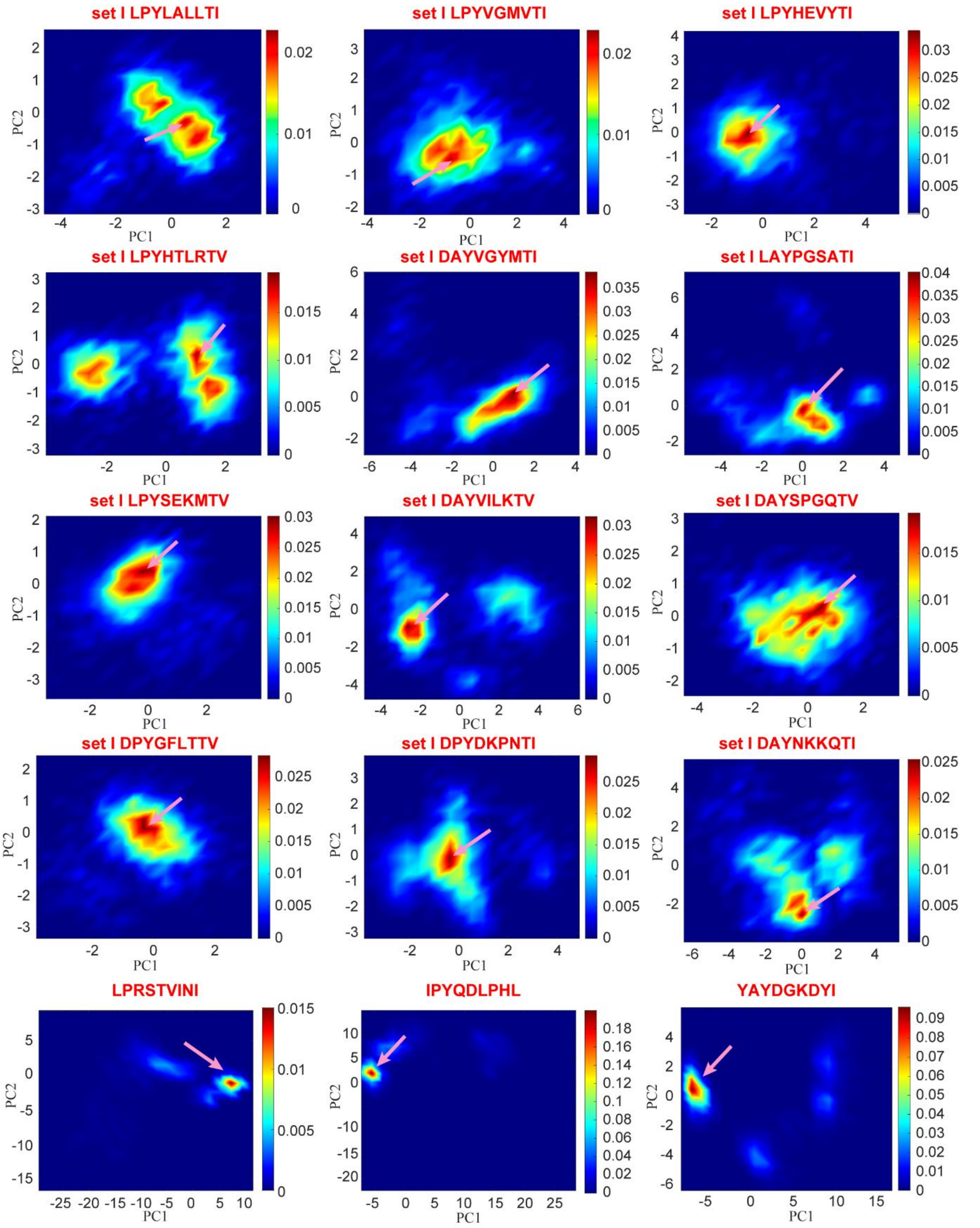


**Figure S1. Distributions of 9-mers binding conformations sampled during conventional molecular dynamics (cMD) simulations.** Two-dimensional conformational density plots are shown as projections of the peptide ensembles onto the first two principal components (PC1 and PC2), after structural alignment of MD frames via the HLA-B*51:01 binding groove. Density maps are generated separately for each peptide in set I. Color scales indicate normalized conformational probability density, and the pink arrow marks the most densely populated binding mode.

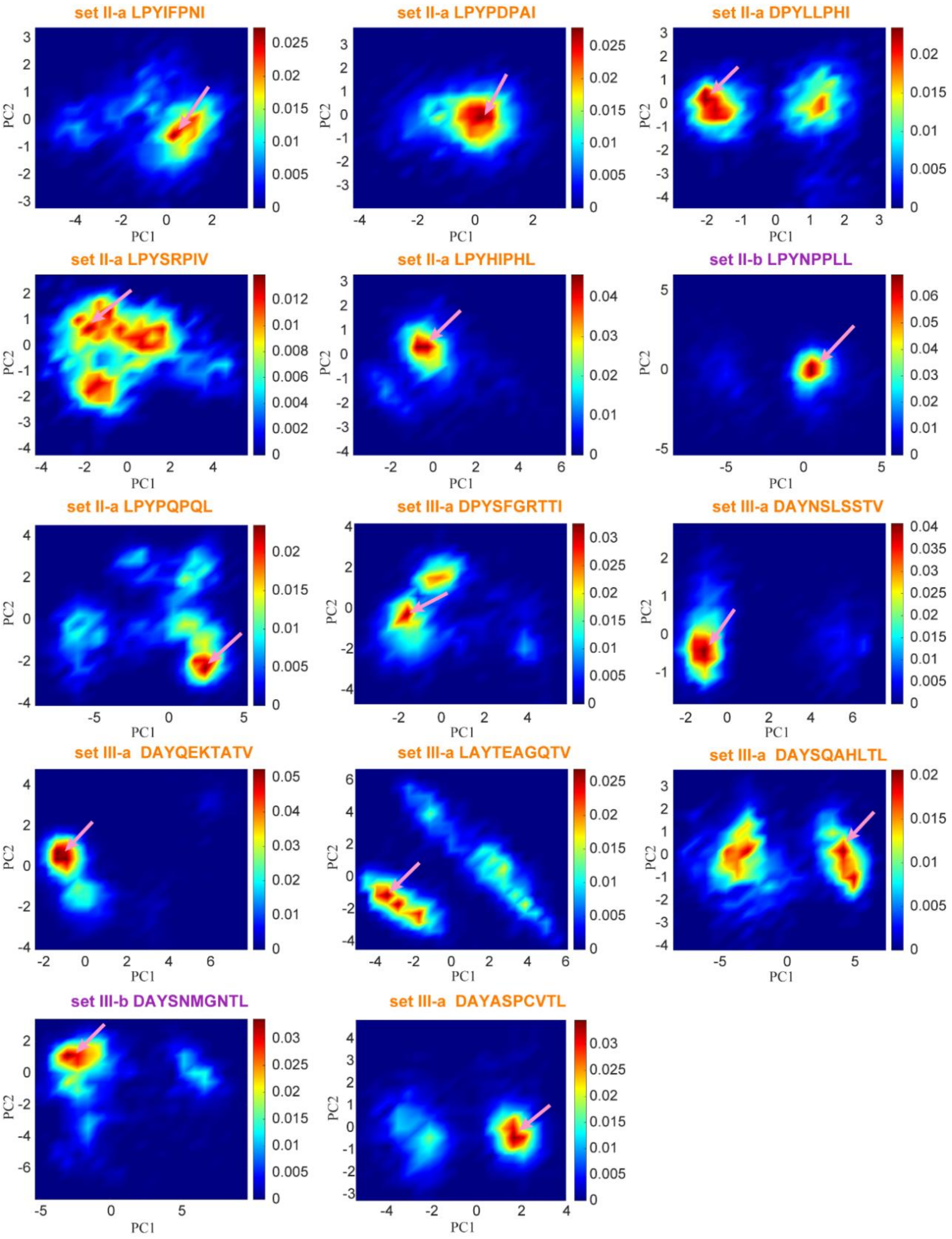


**Figure S2. Distributions of 8-mers and 10-mers binding conformations sampled during cMD simulations.** Two-dimensional conformational density plots are shown as projections of the peptide ensembles onto the first two principal components (PC1 and PC2), after structural alignment of MD frames via the HLA-B*51:01 binding groove. Density maps are generated separately for each peptide in sets II and III. Color scales indicate normalized conformational probability density, and the pink arrow marks the most densely populated binding mode.

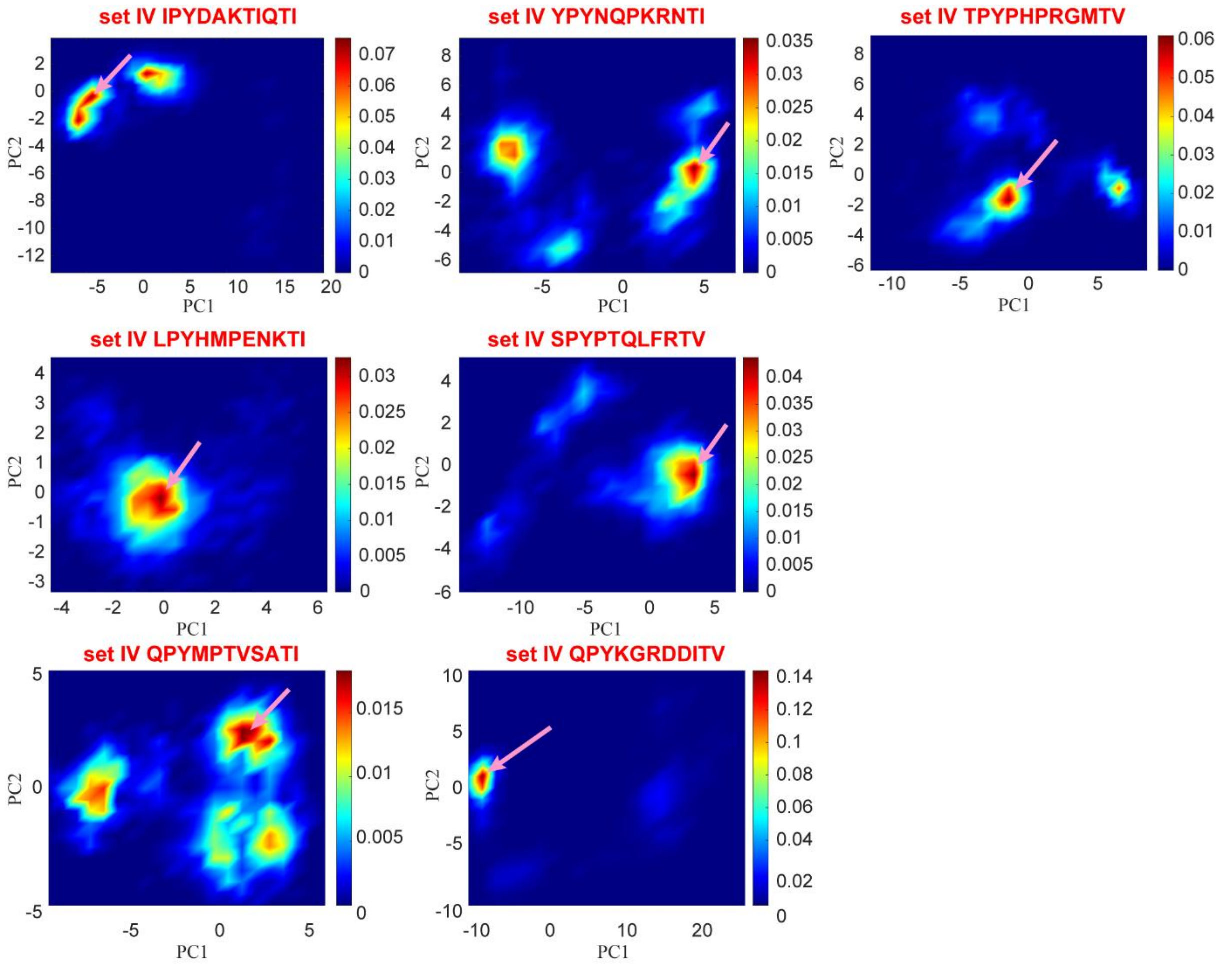


**Figure S3. Distributions of 11-mers binding conformations sampled during cMD simulations.** Two-dimensional conformational density plots are shown as projections of the peptide ensembles onto the first two principal components (PC1 and PC2), after structural alignment of MD frames via the HLA-B*51:01 binding groove. Density maps are generated separately for each peptide in set IV. Color scales indicate normalized conformational probability density, and the pink arrow marks the most densely populated binding mode.

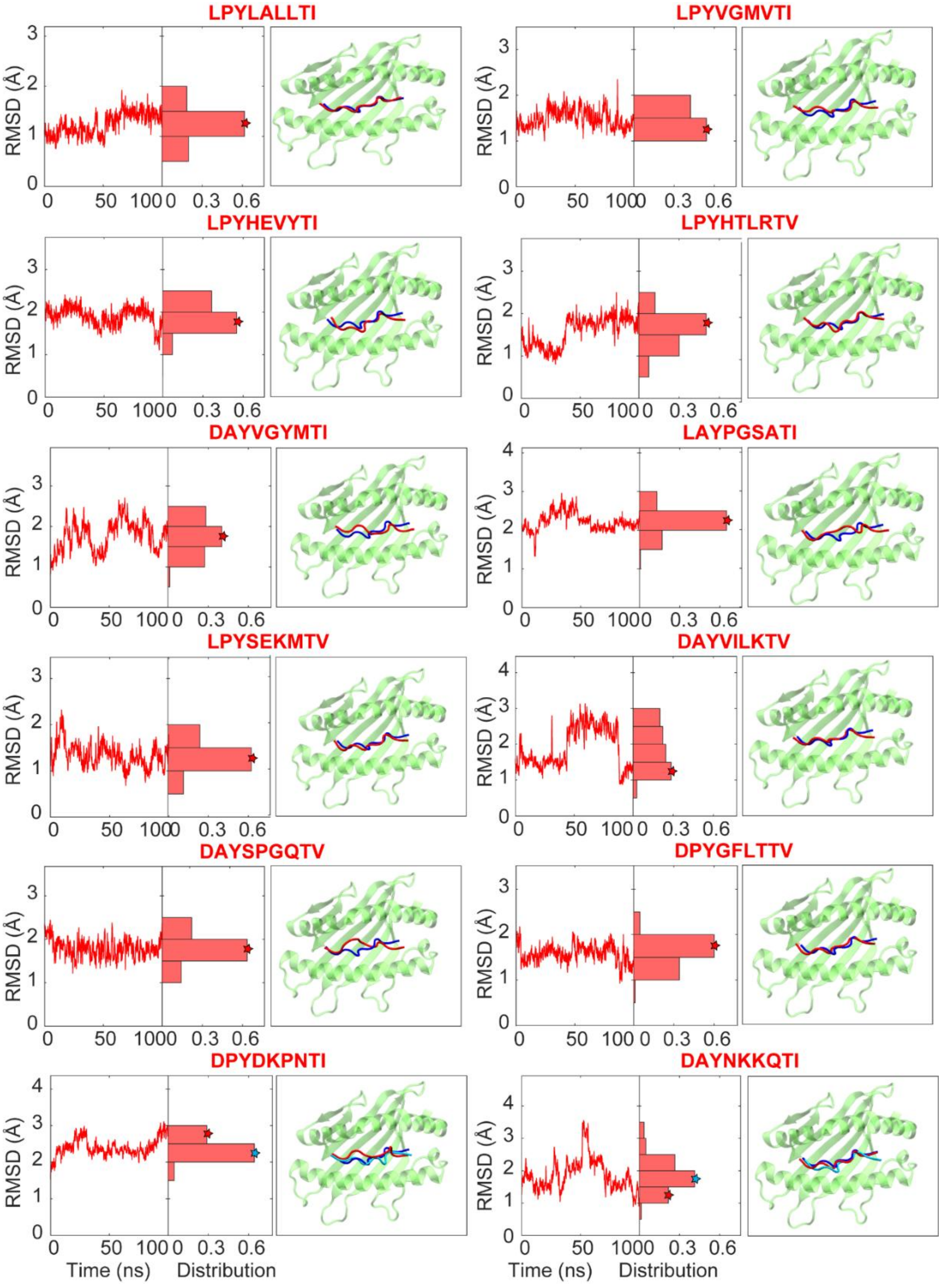


**Figure S4. Root-mean-square deviation (RMSD)-time and RMSD-distribution plots for cMD simulations performed on 9-mers.** In the left panels, the RMSD of each peptide with respect to its starting pose as a function of simulation time, along with the corresponding RMSD distributions, is shown for set I. The red stars indicate the RMSD value of the end conformation within the distribution. In cases where the final conformation does not coincide with the main distribution peak, the highest peak is marked with a cyan star. In the right panels, the HLA-B*51:01 peptide-binding groove (residues G1–Q180) is shown in green. The peptide structures at the end of the trajectories are shown in red. When the most populated RMSD distribution peak does not overlap with the end conformation, a representative peptide conformation corresponding to that peak is additionally shown in cyan. The peptide (LPPVVAKEI) in the crystal structure with PDB: 1E27 is shown in blue.

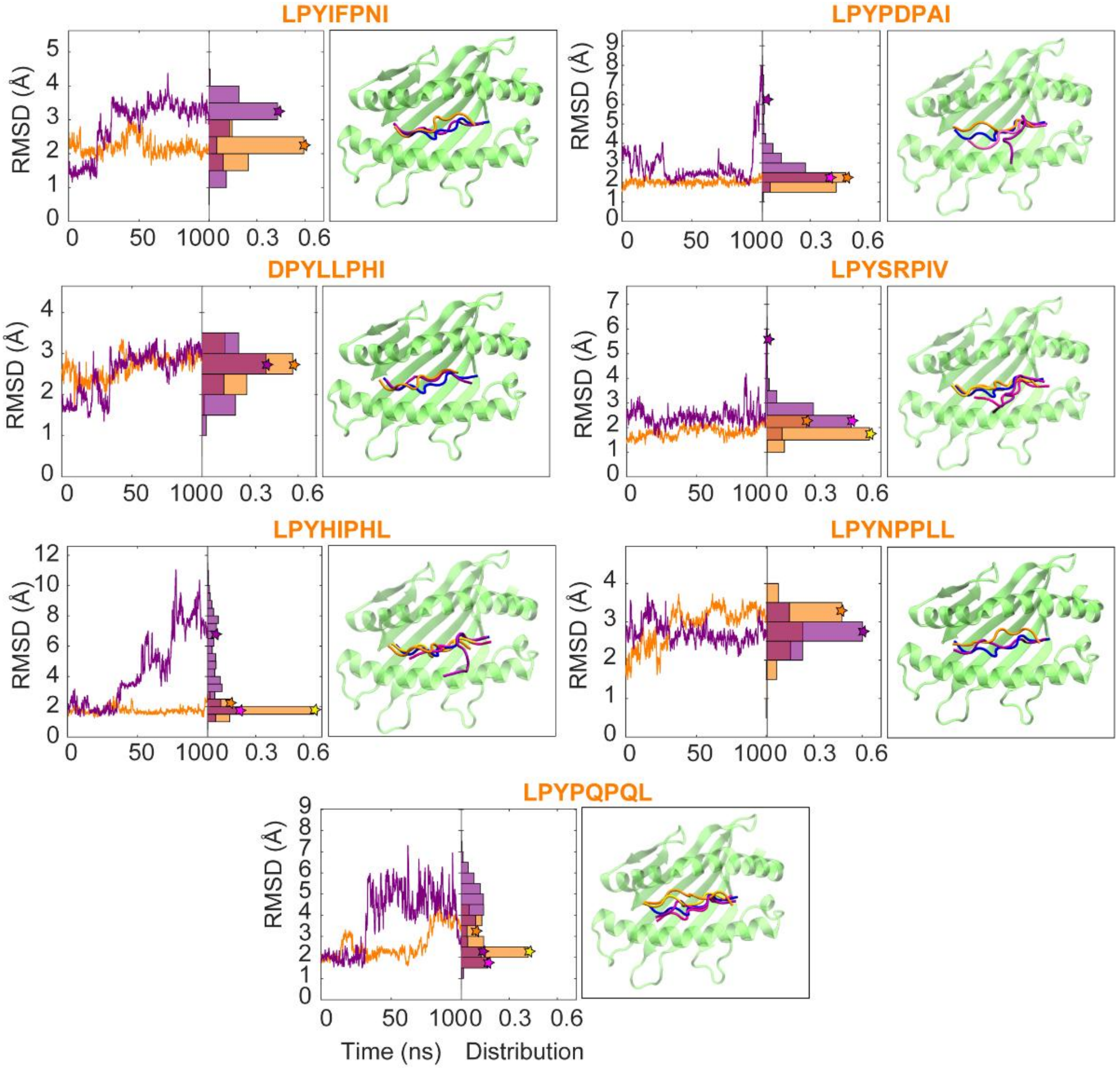


**Figure S5. RMSD-time and RMSD-distribution plots for cMD simulations performed on 8-mers.** In the left panels, the RMSDs of each peptide with respect to its starting pose as a function of simulation time, along with the corresponding RMSD distributions, are shown in orange and purple for sets II-a and II-b, respectively. The orange and purple stars indicate the RMSD values of the end conformations within the distributions. In cases where the final conformations do not coincide with the main distribution peaks, the highest peaks are marked with yellow and violet. In the right panels, the HLA-B*51:01 peptide-binding groove (residues G1–Q180) is shown in green. The peptide structures at the end of the trajectories are shown in orange and purple. When the most populated RMSD distribution peaks do not overlap with the end conformations, representative peptide conformations corresponding to the peaks are additionally shown in yellow and violet. The peptide (LPPVVAKEI) in the crystal structure with PDB: 1E27 is shown in blue.

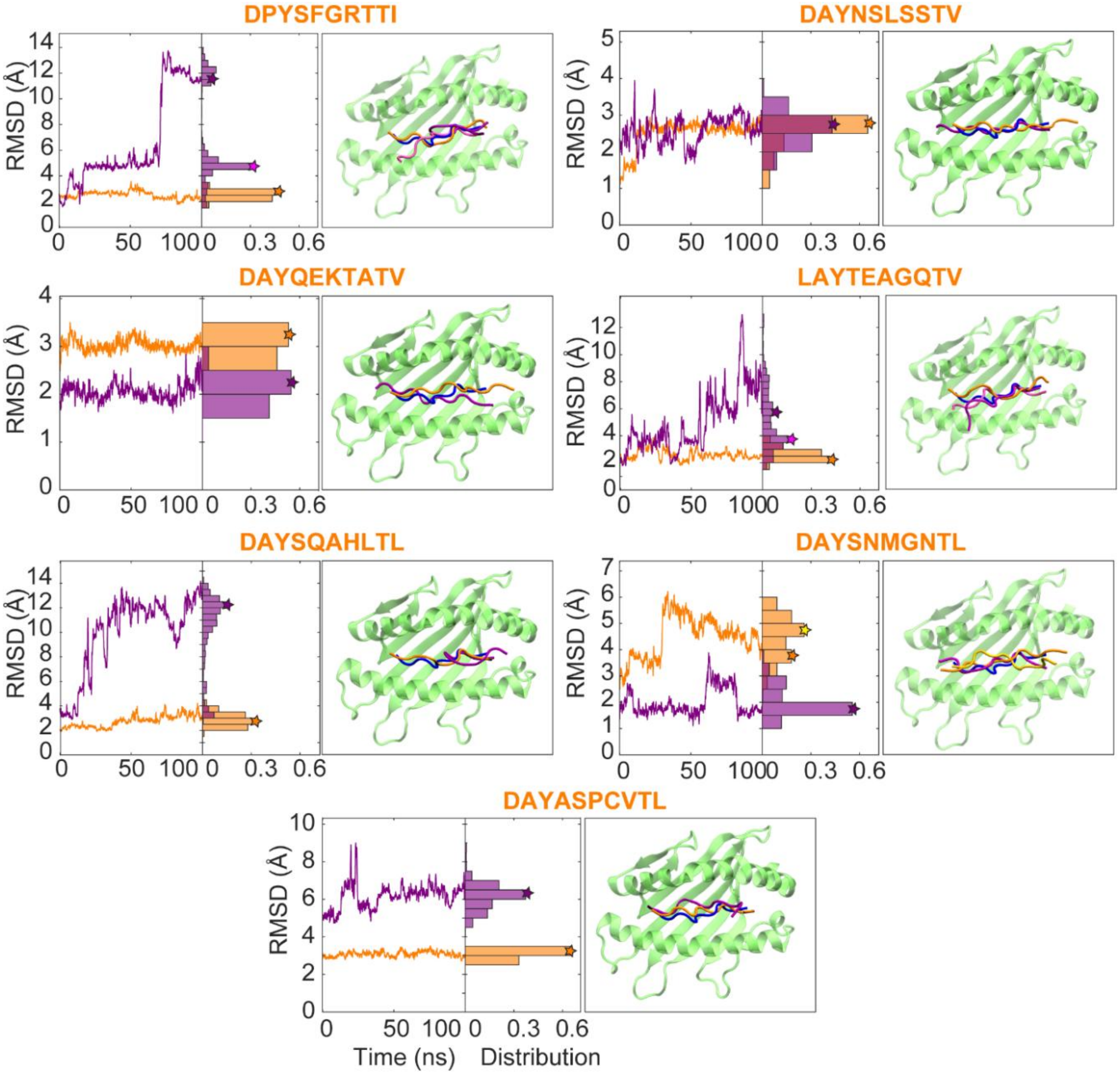


**Figure S6. RMSD-time and RMSD-distribution plots for cMD simulations performed on 10-mers.** In the left panels, the RMSDs of each peptide with respect to its starting pose as a function of simulation time, along with the corresponding RMSD distributions, are shown in orange and purple for sets III-a and III-b, respectively. The orange and purple stars indicate the RMSD values of the end conformations within the distributions. In cases where the final conformations do not coincide with the main distribution peaks, the highest peaks are marked with yellow and violet. In the right panels, the HLA-B*51:01 peptide-binding groove (residues G1–Q180) is shown in green. The peptide structures at the end of the trajectories are shown in orange and purple. When the most populated RMSD distribution peaks do not overlap with the end conformations, representative peptide conformations corresponding to the peaks are additionally shown in yellow and violet. The peptide (LPPVVAKEI) in the crystal structure with PDB: 1E27 is shown in blue.

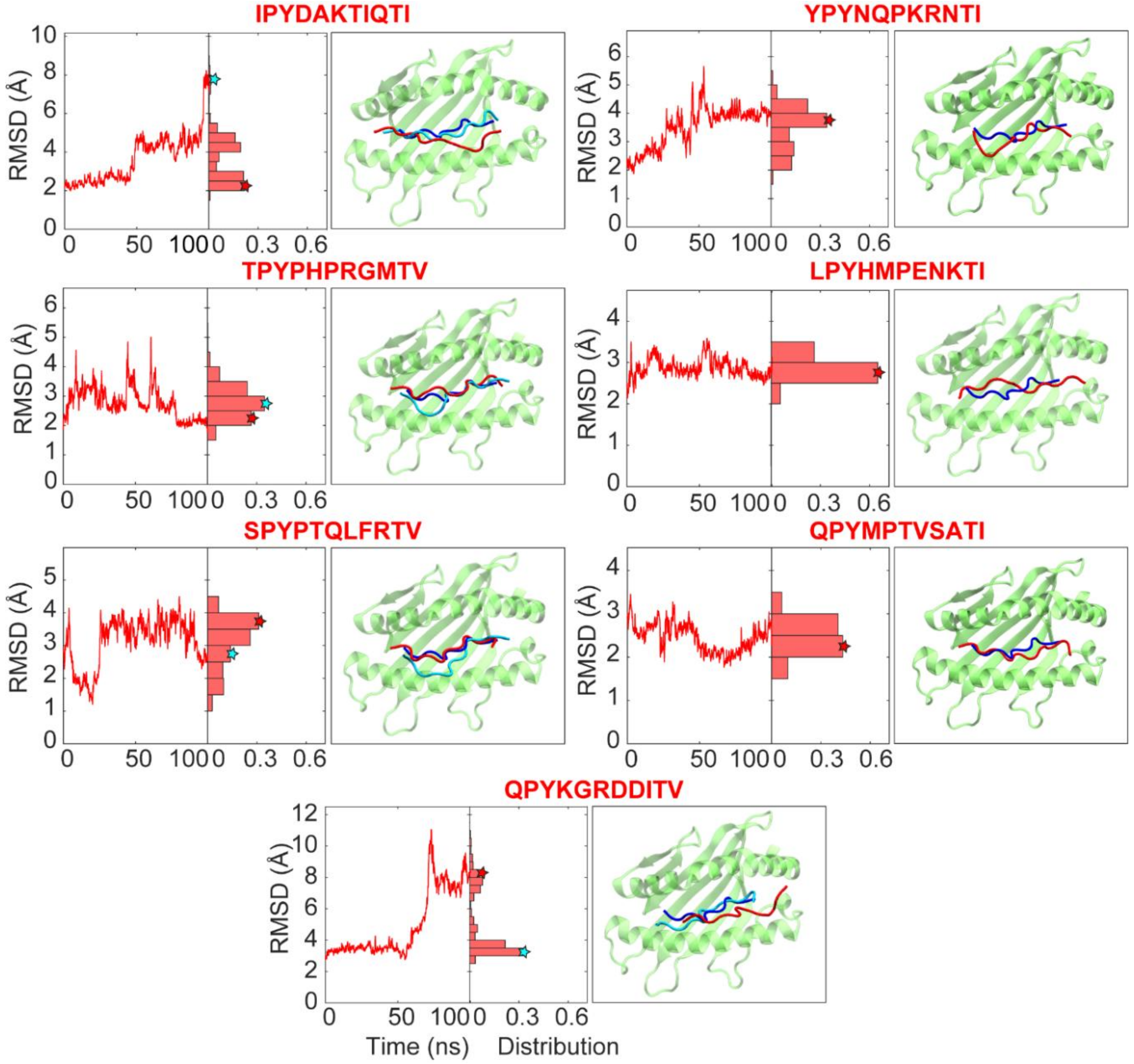


**Figure S7. RMSD-time and RMSD-distribution plots for cMD simulations performed on 11-mers.** In the left panels, the RMSD of each peptide with respect to its starting pose as a function of simulation time, along with the corresponding RMSD distributions, is shown for set VI. The red stars indicate the RMSD value of the end conformation within the distribution. In cases where the final conformation does not coincide with the main distribution peak, the highest peak is marked with a cyan star. In the right panels, the HLA-B*51:01 peptide-binding groove (residues G1–Q180) is shown in green. The peptide structures at the end of the trajectories are shown in red. When the most populated RMSD distribution peak does not overlap with the end conformation, a representative peptide conformation corresponding to that peak is additionally shown in cyan. The peptide (LPPVVAKEI) in the crystal structure with PDB: 1E27 is shown in blue.

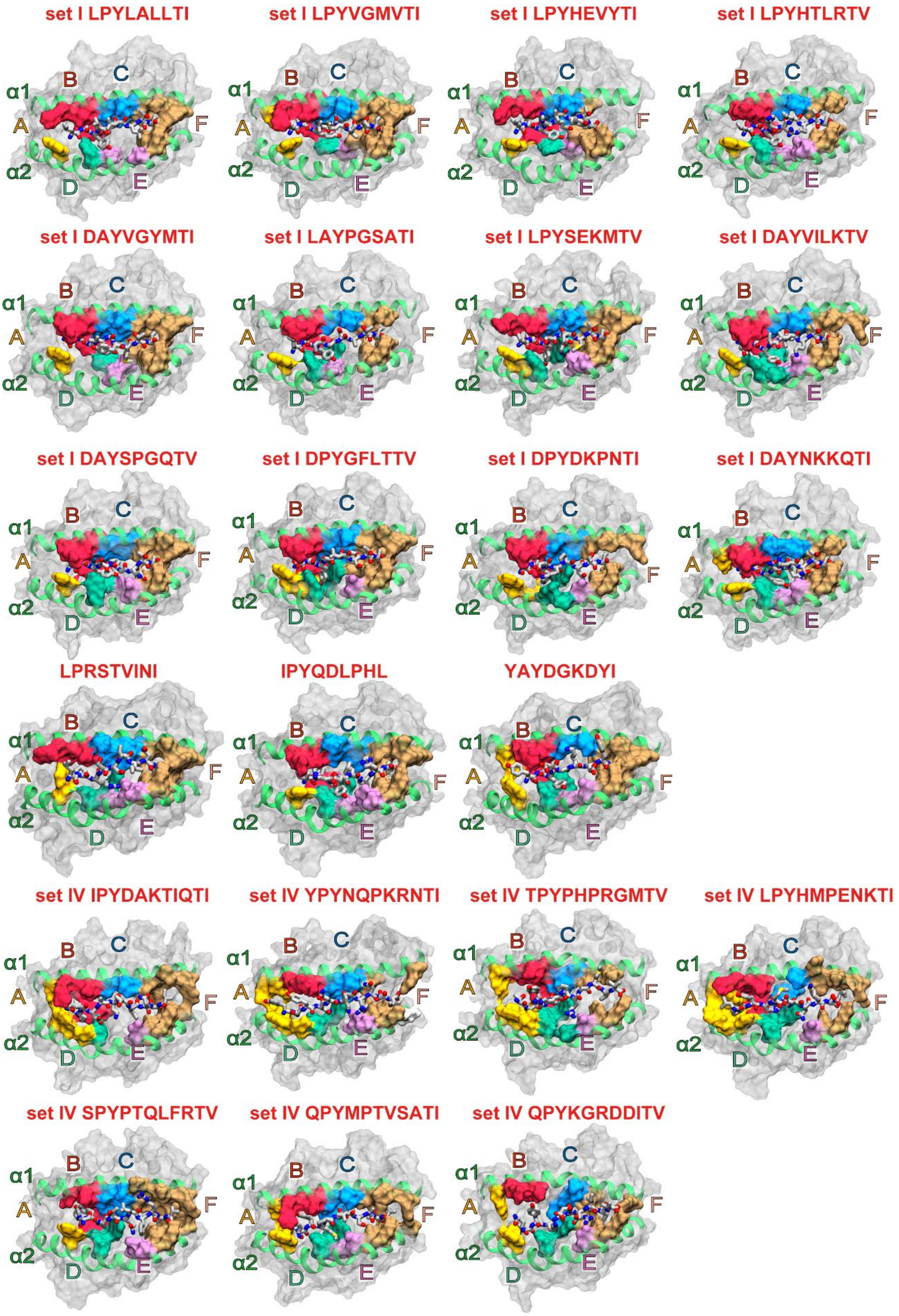


**Figure S8. Representative binding conformations of 9-mers and 11-mers, corresponding to the most densely populated binding mode.** Peptides are shown in licorice representation, together with the binding pockets (A–F) of HLA-B*51:01 are highlighted on the molecular surface (surface representation). Each conformation represents the dominant conformational basin observed in the corresponding density maps shown in Figures S1 and S3.

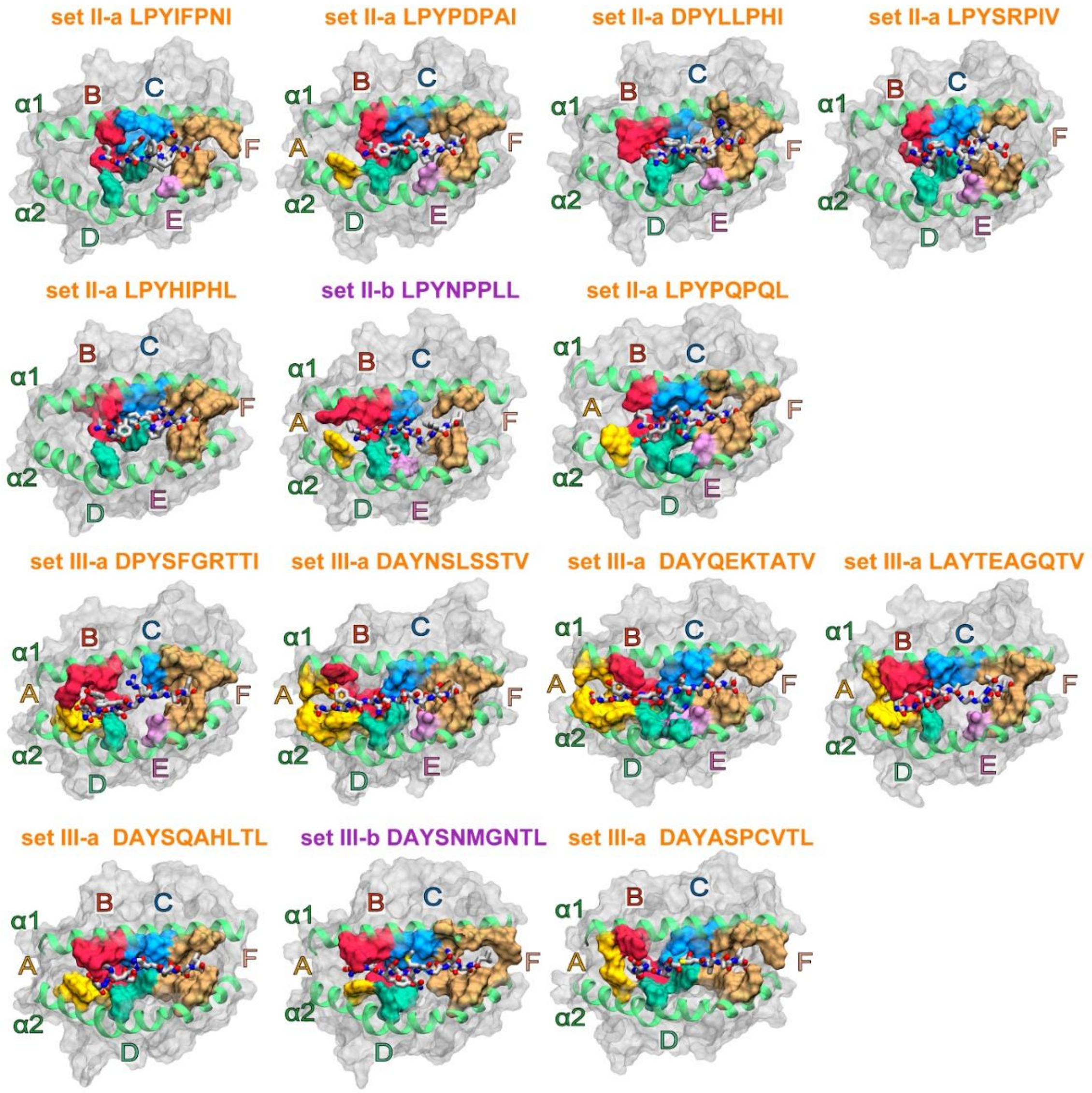


**Figure S9. Representative binding conformations of 8-mers and 10-mers, corresponding to the most densely populated binding mode.** Peptides are shown in licorice representation, while binding pockets (A–F) of HLA-B*51:01 are highlighted on the molecular surface. Each conformation represents the dominant conformational basin observed in the corresponding density maps shown in Figure S2.

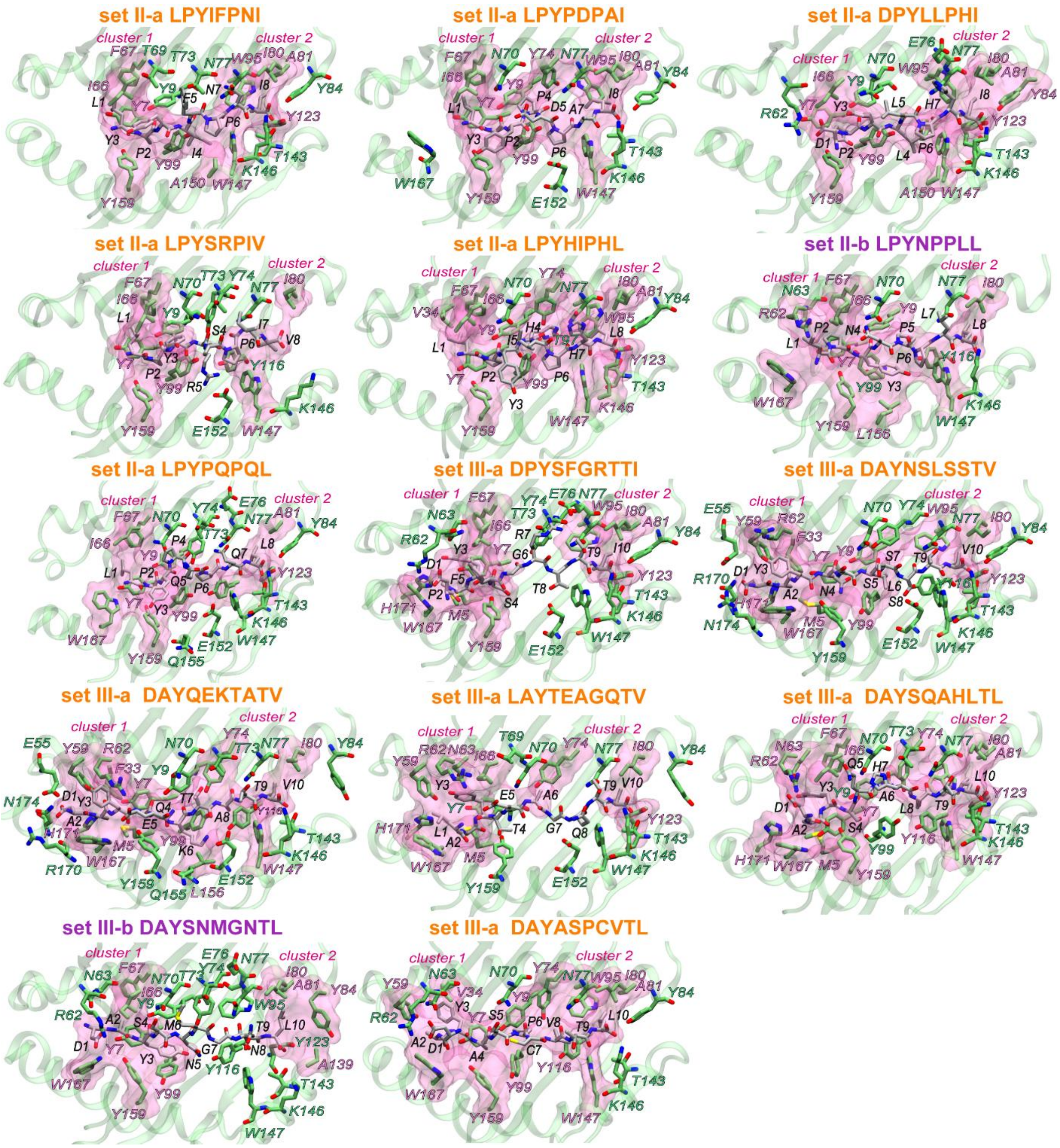


**Figure S10. Two hydrophobic clusters consistently emerge at the interface of HLA-B*51:01 with 8-mers and 10-mers, stabilized by a dense hydrogen bonding network.** Hydrophobic clusters are depicted as transparent magenta surface balloons. Peptide and interacting HLA-B*51:01 residues are shown in licorice representation, with the overall HLA-B*51:01 binding groove rendered in green cartoon representation. Peptide residues are labeled in black. HLA-B*51:01 residues forming hydrophobic contacts with the peptide are labeled in magenta, whereas residues involved in other non-covalent interactions (hydrogen bonds, salt bridges, electrostatic, or π–sulfur interactions) are labeled in green.

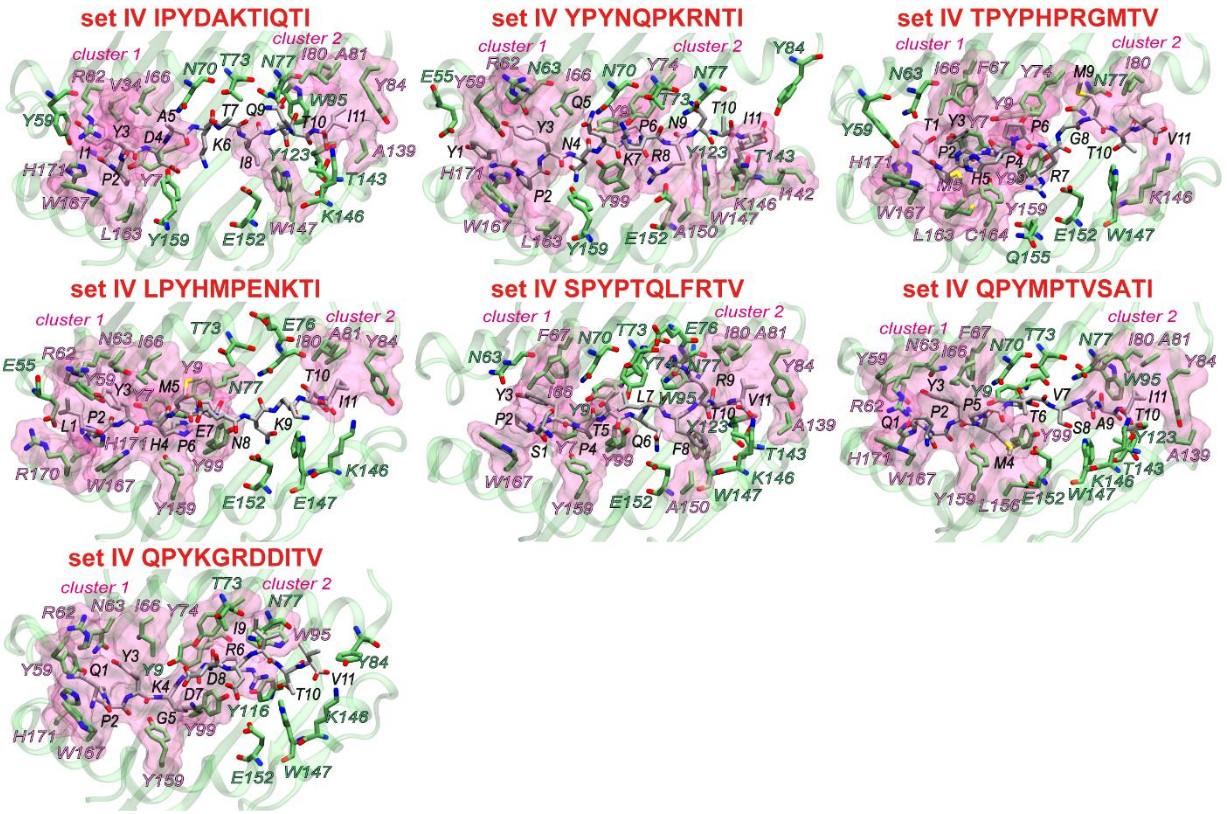


**Figure S11. Two hydrophobic clusters consistently emerge at the interface of HLA-B*51:01 and 11-mers, stabilized by a dense hydrogen bonding network.** Hydrophobic clusters are depicted as transparent magenta surface balloons. Peptide and interacting HLA-B*51:01 residues are shown in licorice representation, with the overall HLA-B*51:01 binding groove rendered in green cartoon representation. Peptide residues are labeled in black. HLA-B*51:01 residues forming hydrophobic contacts with the peptide are labeled in magenta, whereas residues involved in other non-covalent interactions (hydrogen bonds, salt bridges, electrostatic, or π–sulfur interactions) are labeled in green.

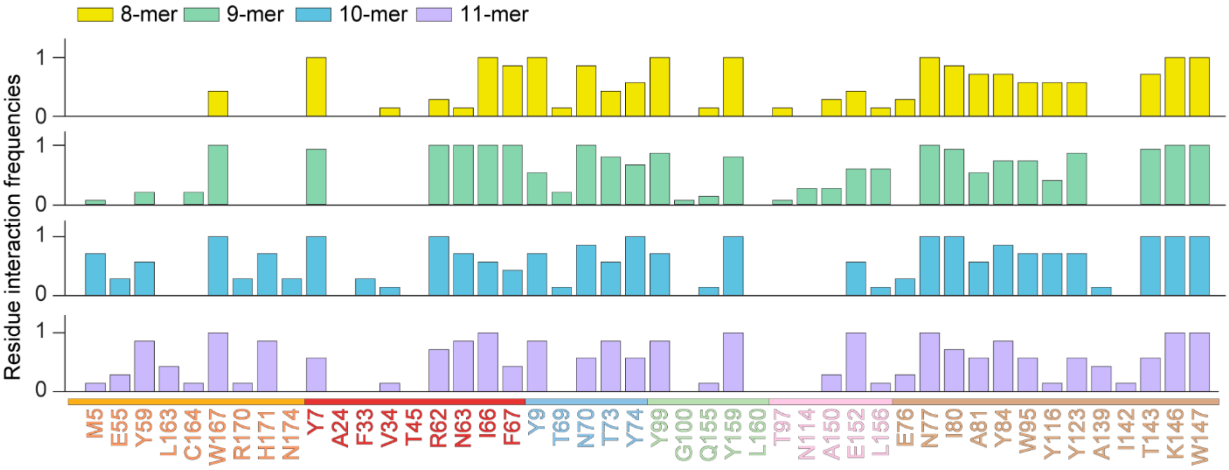


**Figure S12. Length-dependent interaction patterns of peptides bound to HLA-B*51:01**. For every HLA-B*51:01 residue that lines the groove, the mean number of contacts established by peptides of four different lengths (8-, 9-, 10-, and 11-mers). For a given length, contact counts from all peptides were pooled and then divided by the number of peptides in that set, so the resulting bars report the average interactions contributed by each residue and directly expose length-specific preferences.

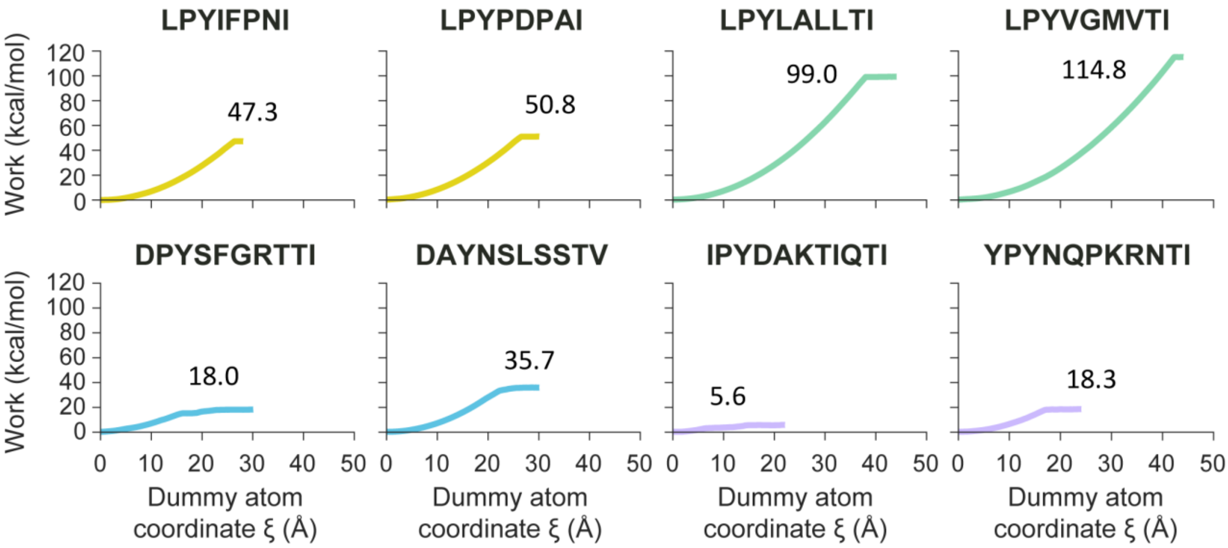


**Figure S13. Work profiles for peptide unbinding from HLA-B*51:01 obtained from steered MD (SMD) simulations.** Work profiles for peptide unbinding from HLA-B*51:01 obtained from SMD simulations. Each plot shows the accumulated work (kcal/mol) as a function of the pulling coordinate (ξ), defined as the distance between the center of mass of the backbone atoms of residue 97 of HLA-B*51:01 and the center of mass of the backbone atoms of the steered peptide residue. The numerical values indicate the total work required for complete peptide dissociation.

#### Supplementary Tables

**Table S1. HLA-B*51:01 binding affinities for peptides in the filtered peptide library, as predicted by NetMHCpan 4.1.**

| No | Peptide | Affinity (nM) | Organism | Source |
|---|---|---|---|---|
| 1 | LPYIFPNI | 67.9 | Human | Ral GTPase-activating protein subunit alpha-1 |
| 2 | LPYPDPAI | 111.3 | Human | Heat shock 70 kDa protein 4 |
| 3 | DPYLLPHI | 215.3 | Human | Serine/threonine-protein kinase tousled-like 2 |
| 4 | LPYSRPIV | 216.3 | Human | Acyl-CoA wax alcohol acyltransferase 1 |
| 5 | LPYHIPHL | 226.3 | Human | E3 ubiquitin-protein ligase listerin |
| 6 | LPYNPPLL | 240.5 | - | Unknown protein eluted from human MHC allele |
| 7 | LPYPQPQL | 260.8 | Wheat | Prolamin |
| 8 | LPYNQPGI | 261.7 | Human | Coatomer subunit gamma-2 |
| 9 | LPYLVPKL | 277.8 | Human | Stalled ribosome sensor GCN1 |
| 10 | LPYVAPEL | 319.2 | Human | Serine/threonine-protein kinase Chk1 |
| 11 | LAYDQPTI | 328.0 | Human | Leucine-rich repeat transmembrane neuronal protein 3 |
| 12 | LPYQPPAL | 465.9 | Human | snRNA-activating protein complex subunit 4 |
| 13 | LPYHPPKL | 520.3 | - | Unknown protein eluted from human MHC allele |
| 14 | LPYPQPGL | 521.1 | Human | Volume-regulated anion channel subunit LRRC8B |
| 15 | DAYVLPKL | 656.0 | Human | Small ribosomal subunit protein eS26 |
| 16 | DAYGVPLL | 665.8 | - | Unknown protein eluted from human MHC allele |
| 17 | DAYTAPAL | 1088.5 | Human | Polymerase delta-interacting protein 3 |
| 18 | LPYLALLTI | 28.6 | Human | Cell cycle checkpoint protein RAD17 |
| 19 | LPYVGMVTI | 29.5 | Human | Signal peptidase complex catalytic subunit SEC11C |
| 20 | LPYHEVYTI | 30.0 | Human | C-type mannose receptor 2 |
| 21 | LPYHTLRTV | 43.6 | Human | Cysteinyl leukotriene receptor 2 |
| 22 | DAYVGYMTI | 45.4 | Human | F-box only protein 11 |
| 23 | LAYPGSATI | 59.2 | Human | WD repeat domain phosphoinositide-interacting protein 2 |
| 24 | LPYSEKMTV | 60.6 | Human | Structural maintenance of chromosomes protein 5 |
| 25 | DAYVILKTV | 99.4 | Human | Gelsolin |
| 26 | DAYSPGQTV | 120.2 | Human | Complement C5 |
| 27 | DPYGFLTTV | 129.6 | Human | Small integral membrane protein 15 |
| 28 | DPYDKPNTI | 228.6 | Human | Sortilin-related receptor |
| 29 | DAYNKKQTI | 251.9 | Human | CMP-N-acetylneuraminate-beta-galactosamide-alpha-2,3-sialyltransferase 4 |
| 30 | DPYSFGRTTI | 433.8 | Human | CSC1-like protein 1 |
| 31 | DAYNSLSSTV | 538.7 | Human | Integrin beta-7 |
| 32 | DAYQEKTATV | 932.7 | Human | NADP-dependent malic enzyme |
| 33 | LAYTEAGQTV | 1019.5 | Human | Nucleoporin NUP188 |
| 34 | DAYSQAHLTL | 1131.5 | Human | pre-rRNA 2'-O-ribose RNA methyltransferase FTSJ3 |
| 35 | DAYSNMGNTL | 1761.5 | Human | UDP-N-acetylglucosamine--peptide N-acetylglucosaminyltransferase 110 kDa subunit |
| 36 | DAYASPCVTL | 1854.0 | - | Unknown protein eluted from human MHC allele |
| 37 | IPYDAKTIQTI | 220.1 | Human | Leucine-rich repeat and coiled-coil domain-containing protein 1 |
| 38 | YPYNQPKRNTI | 326.3 | Human | RNA helicase aquarius |
| 39 | TPYPHPRGMTV | 391.9 | Human | Postmeiotic segregation increased 2-like protein 5 |
| 40 | LPYHMPENKTI | 401.0 | Human | Claspin |
| 41 | SPYPTQLFRTV | 857.0 | Human | ETS-related transcription factor Elf-1 |
| 42 | QPYMPTVSATI | 1405.4 | Human | Methionine--tRNA ligase, cytoplasmic |
| 43 | QPYKGRDDITV | 6686.3 | Human | Glycoprotein endo-alpha-1,2-mannosidase-like protein |

##### Table S2. cMD simulations of peptides binding to HLA-B*51:01.

| Set | Template pose | Length | Bound peptide | Duration (ns) |
|---|---|---|---|---|
| I | LPPVVAKEI (P1-P9) | 9 | LPYLALLTI | 100 |
| | | | LPYVGMVTI | 100 |
| | | | LPYHEVYTI | 100 |
| | | | LPYHTLRTV | 100 |
| | | | DAYVGYMTI | 100 |
| | | | LAYPGSATI | 100 |
| | | | LPYSEKMTV | 100 |
| | | | DAYVILKTV | 100 |
| | | | DAYSPGQTV | 100 |
| | | | DPYGFLTTV | 100 |
| | | | DPYDKPNTI | 100 |
| | | | DAYNKKQTI | 100 |
| II | a) -PPVVAKEI (P2-P9)<br>b) LPPVVAKE- (P1-P8) | 8 | LPYIFPNI | a) 100, b) 100 |
| | | | LPYPDPAI | a) 100, b) 100 |
| | | | DPYLLPHI | a) 100, b) 100 |
| | | | LPYSRPIV | a) 100, b) 100 |
| | | | LPYHIPHL | a) 100, b) 100 |
| | | | LPYNPPLL | a) 100, b) 100 |
| | | | LPYPQPQL | a) 100, b) 100 |
| III | a) +LPPVVAKEI (Adding at N-terminal),<br>b) LPPVVAKEI+ (Adding at C-terminal) | 10 | DPYSFGRTTI | a) 100, b) 100 |
| | | | DAYNSLSSTV | a) 100, b) 100 |
| | | | DAYQEKTATV | a) 100, b) 100 |
| | | | LAYTEAGQTV | a) 100, b) 100 |
| | | | DAYSQAHLTL | a) 100, b) 100 |
| | | | DAYSNMGNTL | a) 100, b) 100 |
| | | | DAYASPCVTL | a) 100, b) 100 |
| VI | +LPPVVAKEI+ (Adding at N- and C-terminal) | 11 | IPYDAKTIQTI | 100 |
| | | | YPYNQPKRNTI | 100 |
| | | | TPYPHPRGMTV | 100 |
| | | | LPYHMPENKTI | 100 |
| | | | SPYPTQLFRTV | 100 |
| | | | QPYMPTVSATI | 100 |
| | | | QPYKGRDDITV | 100 |

*Locations of residue additions (+) and deletions (–) relative to the reference peptide LPPVVAKEI, which was used to model peptides in the binding cavity, are highlighted in orange.*

##### Table S3. SMD simulations of peptides pulled from HLA-B*51:01.

| Set | Template pose | Length | Bound peptide | Duration (ns) |
|---|---|---|---|---|
| **I** | LPPVVAKEI (P1-P9) | 9 | LPYLALLTI<br>LPYVGMVTI | 440<br>440 |
| **II** | a) –LPPVVAKE (P2-P9) | 8 | LPYIFPNI<br>LPYPDPAI | 280<br>300 |
| **III** | a) +LPPVVAKEI (Adding at N-terminal) | 10 | DPYSFGRTTI<br>DAYNSLSSTV | 300<br>300 |
| **VI** | +LPPVVAKEI+ (Adding at N- and C-terminal) | 11 | IPYDAKTIQTI<br>YPYNQPKRNTI | 220<br>240 |

*Locations of residue additions (+) and deletions (–) relative to the reference peptide LPPVVAKEI, which was used to model peptides in the binding cavity, are highlighted in orange.*

#### Table S4. Binding motifs of 8-mers to HLA-B*51:01.

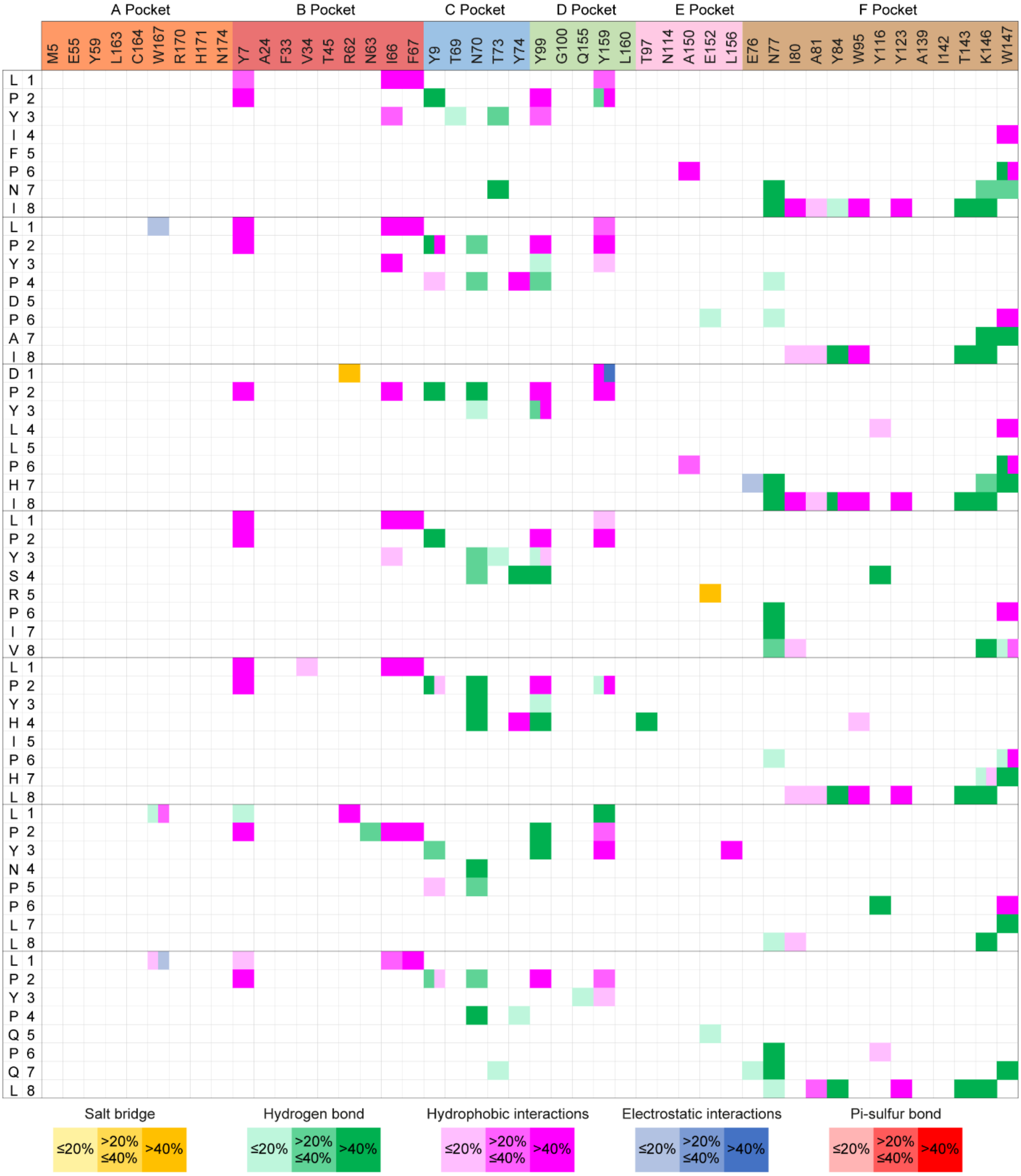

#### Table S5. Binding motifs of 9-mers to HLA-B*51:01.

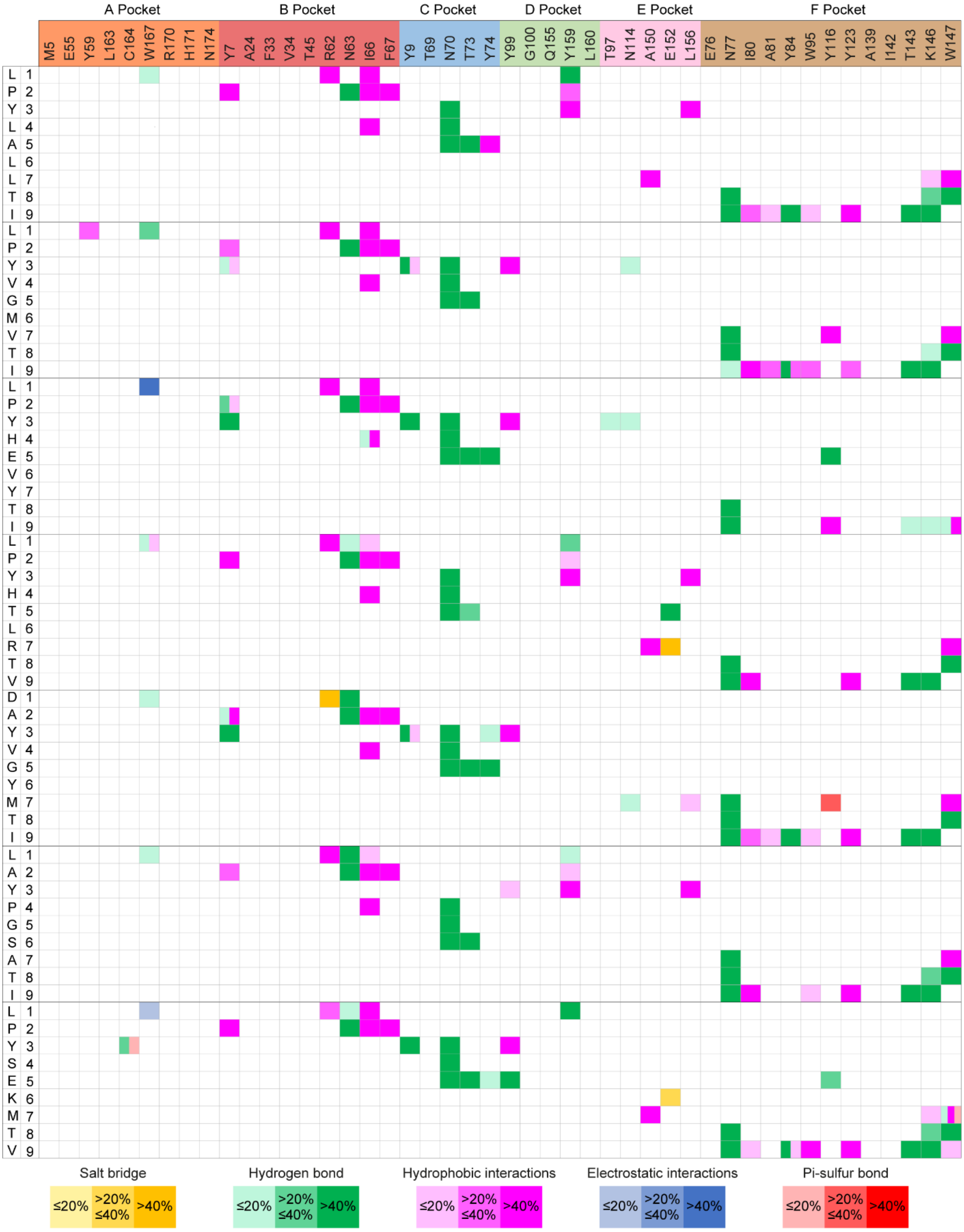

#### Table S6. Binding motifs of 9-mers to HLA-B*51:01.

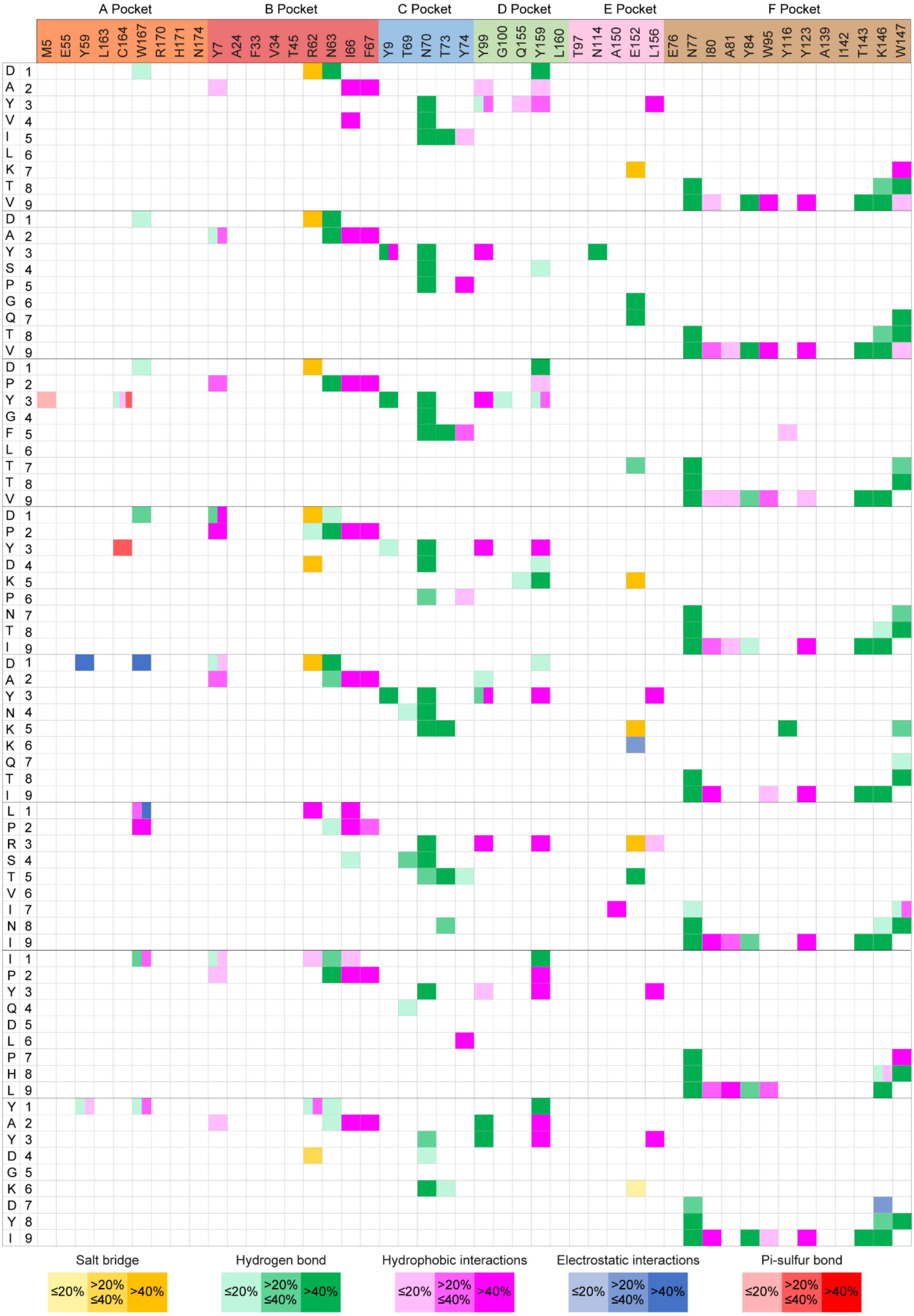

#### Table S7. Binding motifs of 10-mers to HLA-B*51:01.

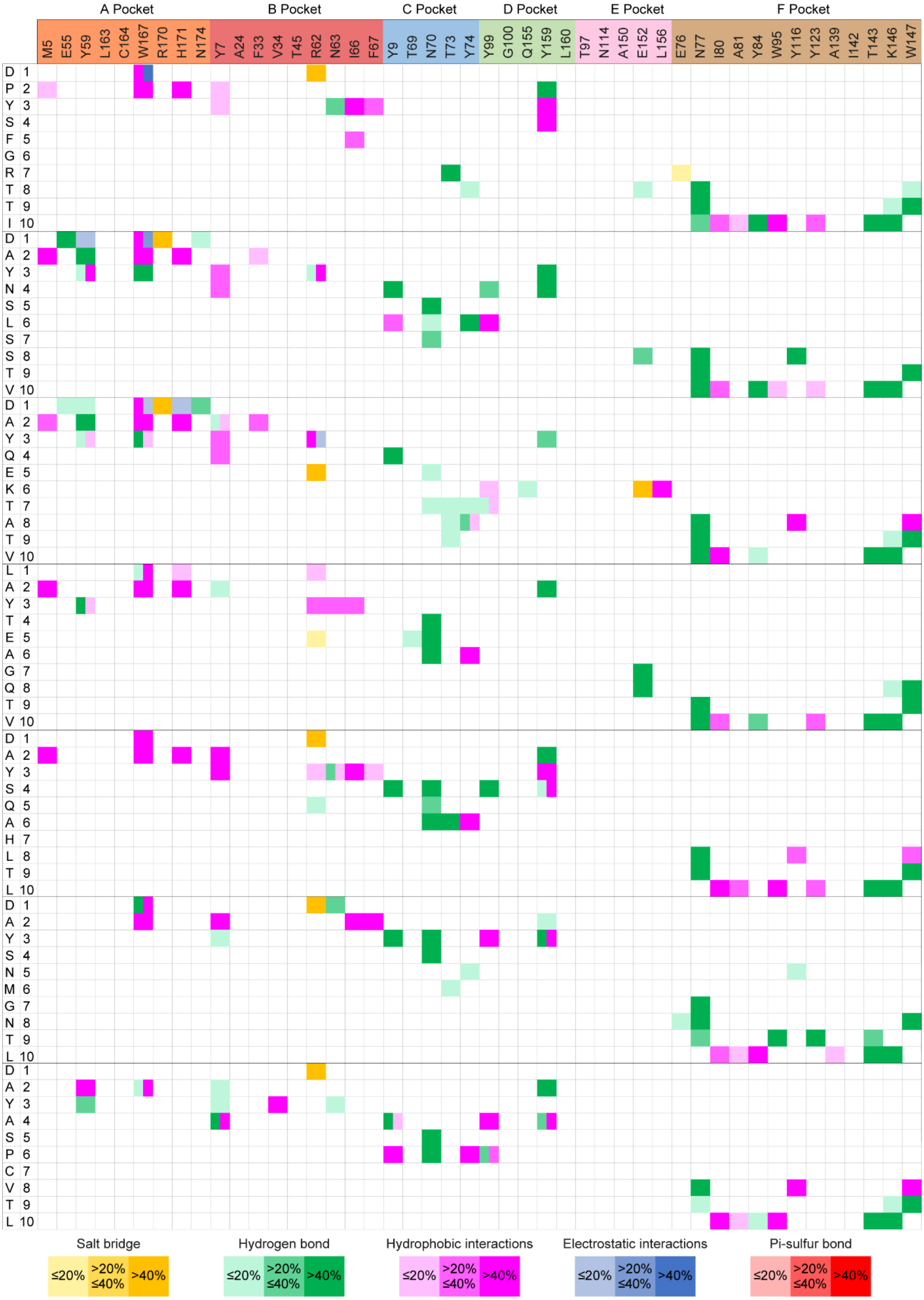

#### Table S8. Binding motifs of 11-mers to HLA-B*51:01.

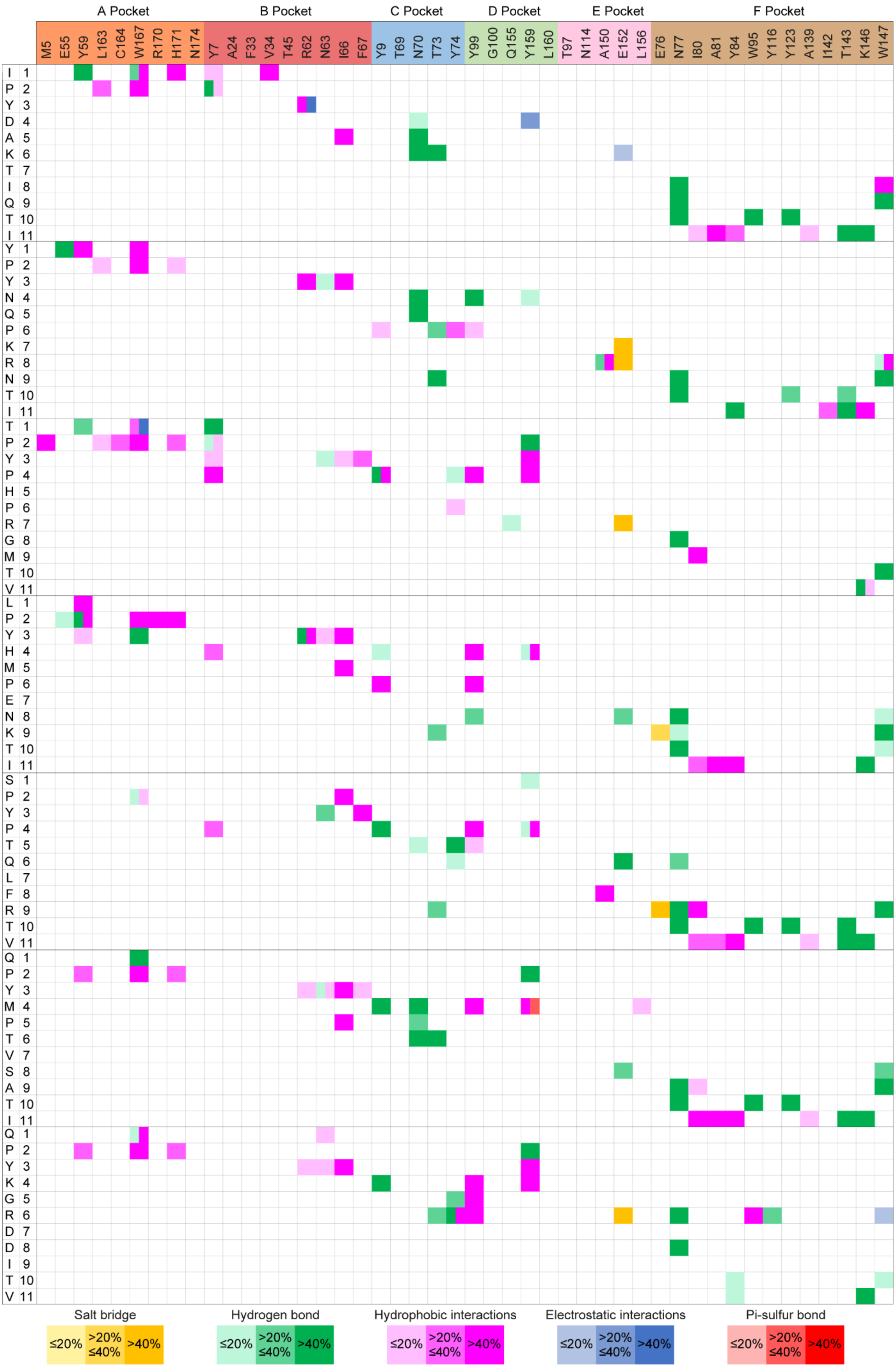
